# Within-host evolutionary dynamics of seasonal and pandemic human influenza A viruses in young children

**DOI:** 10.1101/2021.03.20.436248

**Authors:** Alvin X. Han, Zandra C. Felix Garza, Matthijs R. A. Welkers, René M. Vigeveno, Tran Nhu Duong, Le Thi Quynh Mai, Pham Quang Thai, Dang Dinh Thoang, Tran Thi Ngoc Anh, Ha Manh Tuan, Nguyen Thanh Hung, Le Quoc Thinh, Le Thanh Hai, Hoang Thi Bich Ngoc, Kulkanya Chokephaibulkit, Pilaipan Puthavathana, Nguyen Van Vinh Chau, Nghiem My Ngoc, Nguyen Van Kinh, Dao Tuyet Trinh, Tran Tinh Hien, Heiman F. L. Wertheim, Peter Horby, Annette Fox, H. Rogier van Doorn, Dirk Eggink, Menno D. de Jong, Colin A. Russell

## Abstract

The evolution of influenza viruses is fundamentally shaped by within-host processes. However, the within-host evolutionary dynamics of influenza viruses remain incompletely understood, in part because most studies have focused on within-host virus diversity of infections in otherwise healthy adults based on single timepoint data. Here, we analysed the within-host evolution of 82 longitudinally-sampled individuals, mostly young children, infected with A/H3N2 or A/H1N1pdm09 viruses between 2007 and 2009. For A/H1N1pdm09 infections during the 2009 pandemic, nonsynonymous changes were common early in infection but decreased or remained constant throughout infection. For A/H3N2 viruses, early infection was dominated by purifying selection. However, as infections progressed, nonsynonymous variants increased in frequencies even though within-host virus titres decreased, leading to the maintenance of virus diversity via mutation-selection balance. Our findings suggest that this maintenance of genetic diversity in these children combined with their longer duration of infection may provide important opportunities for within-host virus evolution.

## Introduction

Influenza A viruses (IAV) are some of the most prevalent human respiratory pathogens, infecting hundreds of millions of people worldwide each year. Because of the high error rates of the viral RNA polymerase complex, *de novo* mutants are generated as the viruses replicate within infected hosts^1^. However, the emergence of these variants within host does not mean that they will become the majority variant within the infected host or be transmitted between hosts. The evolution of IAVs is the product of a complex mosaic of evolutionary processes that include genetic drift, positive selection^2^, transmission bottleneck effects^3,4^ and global migration patterns^5,6^. Importantly, the resulting evolutionary dynamics can differ at the individual and population levels^7^.

For seasonal IAVs at the global population level, antibody-mediated immune selection pressure from natural infection or vaccination positively selects for novel antigenic variants that facilitate immune escape resulting in antigenic drift^2^. However, at the within-host level, the role of positive selection exerted by immunity is less obvious. Several next generation sequencing studies of typical, short-lived seasonal IAV infections in adult humans showed that intra-host genetic diversity of influenza viruses is low and dominated by purifying selection^4,8–11^. Additionally, large scale comparative analyses of IAV haemagglutinin (HA) consensus sequences found limited evidence of positive selection on HA at the individual level regardless of the person’s expected influenza virus infection history^12^. Importantly, these studies focused on virus samples from only one or two time points, mostly early in infection, limiting the opportunities to study how virus populations evolved over the course of infection.

Separate from seasonal IAVs, zoonotic IAVs constantly pose new pandemic threats. Prior to becoming human-adapted seasonal strains, IAVs are introduced into the human population from an animal reservoir through the acquisition of host adaptive mutations, sometimes via reassortment, resulting in global pandemics such as the 2009 swine influenza pandemic^13^. In the 2009 pandemic, global virus genetic diversity increased rapidly during the early phases of the pandemic as a result of rapid transmissions in the predominantly naïve human population^14^. Over subsequent waves of the pandemic, host adapting mutations that incrementally improved viral fitness and transmissibility in humans of A/H1N1pdm09 viruses emerged^15^, eventually reaching fixation in the global virus population^16^.

At the individual level, the within-host evolutionary dynamics of the pandemic A/H1N1pdm09 virus, particularly in the early stages of the 2009 pandemic, have been relatively underexplored. To date, the only within-host genetic diversity analysis of A/H1N1pdm09 viruses during the initial phase of the pandemic was based on mostly single-timepoint samples collected within ∼7 days post-symptom onset^17^. Despite initial findings of high within-host diversity and loose transmission bottlenecks^17^, these results were later disputed due to technical anomalies and subsequent reanalyses of a smaller subset of the original data found that intra-host genetic diversity of the pandemic virus was low and comparable to levels observed in seasonal IAVs^18,19^. It remains unclear how frequently host adaptive mutations appear within hosts infected by a pandemic IAV and if these mutants are readily transmitted between individuals.

Here, we deep sequenced 275 longitudinal clinical specimens sampled from 82 individuals residing in Southeast Asia between 2007 and 2009 that were either infected with seasonal A/H3N2 or pandemic A/H1N1pdm09 viruses. By analysing minority variants found across the whole IAV genome, we characterised the evolutionary dynamics of within-host virus populations in these samples collected up to two weeks post-symptom onset.

## Results

### Study participants

The A/H3N2 virus samples were collected from 51 unlinked individuals as part of an oseltamivir dosage trial^20,21^. 48 of the 51 A/H3N2 virus infected individuals were young children (median age=2 years; interquartile range (IQR)=2-3 years) at the time of sampling and most had low or no detectable anti-influenza virus antibody titers on day 0 and 10 post-symptom onset^21^. Given that young children are substantial contributors to influenza virus transmission^22,23^, the samples analysed here offer a valuable opportunity to investigate the within-host IAV evolutionary dynamics in this key population. The A/H1N1pdm09 virus specimens were collected from 32 individuals up to 12 days post-symptom onset. These individuals include both children and adults (median age=10 years; IQR=4-20 years) infected during the first wave of the pandemic in Vietnam (July-December 2009). 15 of the 32 individuals (including 6 index patients) were sampled in a household-based influenza cohort study^24^. The remaining 16 unlinked individuals were hospitalised patients that were involved in two different oseltamivir treatment studies^20,25^ Details of all study participants are described in the respective cited studies and Table S4.

### Genetic diversity of within-host virus populations

We used the number of minority intra-host single nucleotide variants (iSNVs; ≥ 2% in frequencies) to measure the levels of genetic diversity of within-host IAV populations. Similar to previous studies^4,8,9,11^, within-host genetic diversity of human A/H3N2 virus populations was low (median = 11 iSNVs, interquartile-range (IQR) = 7-16; Figure 1A). Within-host genetic diversity of pandemic A/H1N1pdm09 virus populations was also low, with a median number of 21 iSNVs (IQR = 13.5-30.0; Figure 1B) identified. Cycle threshold (Ct) values, and thus likely virus shedding, correlated with the number of days post-symptom onset for both IAV subtypes (A/H3N2: Spearman’s *ρ* = 0.468, *p* = 1.38 × 10^−10^; A/H1N1pdm09: *ρ* = 0.341, *p* = 0.048; Figure 1C and D). The number of iSNVs observed in within-host A/H3N2 virus populations weakly correlated with days since onset of symptoms in patients (*ρ* = 0.463, *p* = 2.22 × 10^−10^) and Ct values (ρ = 0.508, *p* = 1.20 × 10^−12^), suggesting that as infection progresses, genetic variants accumulate within-host even as virus population size decreases (Figure 1A). On the other hand, there was no significant correlation between the number of iSNVs observed in within-host A/H1N1pdm09 virus populations and Ct values (*ρ* = 0.1 8, *p* = 0.21) or days post-symptom onset (*ρ* = −0.021, *p* = 0. 1) (Figure 1B).

**Figure 1.**
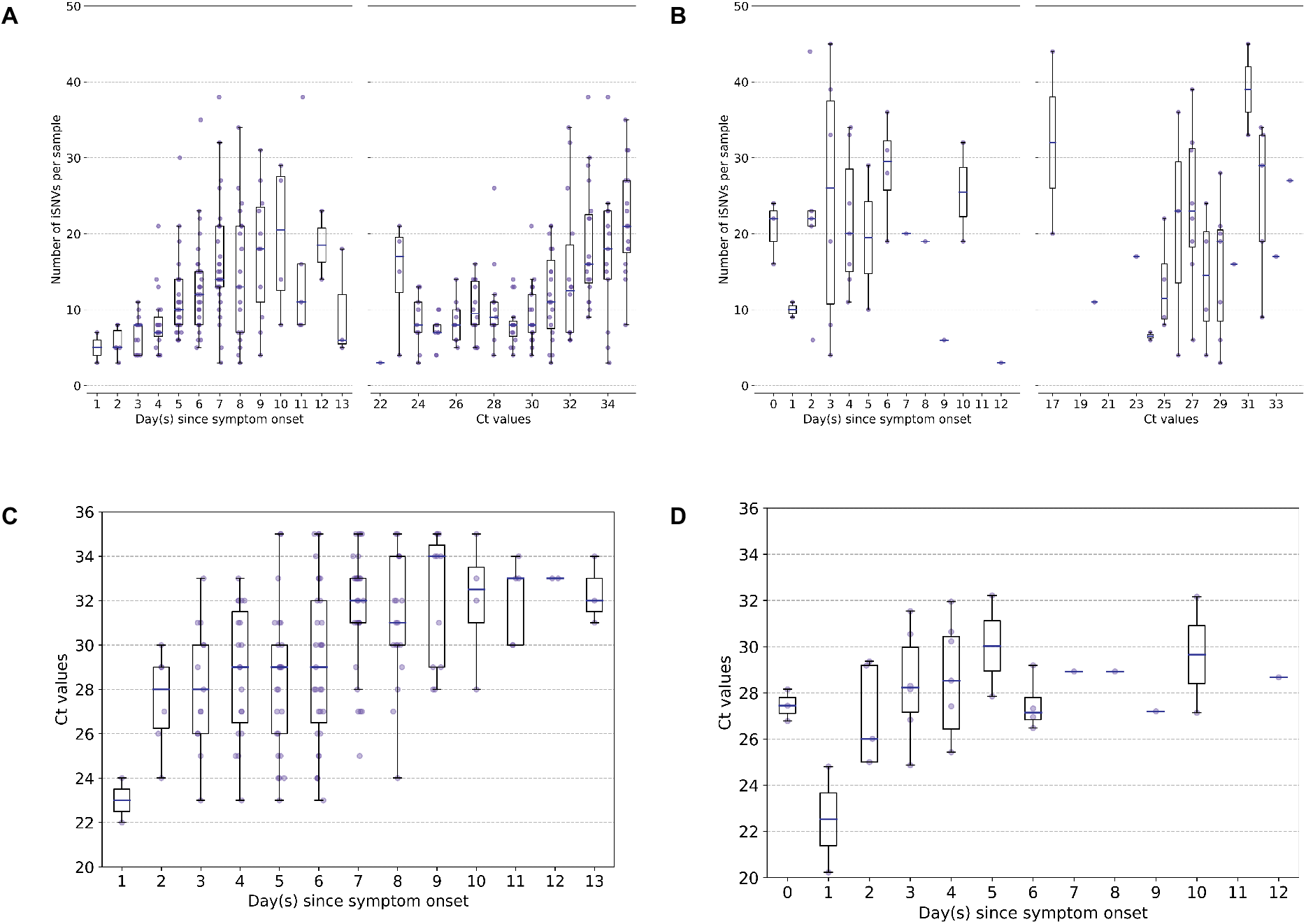
Genetic diversity of within-host influenza A virus populations. Box plots summarizing the number of intra-host single nucleotide variants (iSNVs; median, interquartile range (IQR), and whiskers extending within median ±1.5 × IQR) identified in samples with adequate breadth of coverage across the whole influenza virus genome in (A) seasonal A/H3N2 and (B) pandemic A/H1N1pdm09 virus samples, stratified by day(s) since symptom onset or qPCR cycle threshold (Ct) values. (C, D) Ct values as a function of day(s) since symptom onset for A/H3N2 viruses (C) and A/H1N1pdm09 viruses (D).

### Within-host evolutionary rates of influenza A viruses

To investigate within-host evolutionary dynamics, empirical rates of synonymous, non-synonymous, and premature stop-codon (i.e. nonsense) iSNVs were calculated by normalizing the summation of observed iSNV frequencies with the number of available sites and time since symptom onset (see Methods). The overall within-host evolutionary rates of A/H3N2 viruses observed here are in the same order of magnitude (< ∽10^−5^ divergence per site per day) as those reported in previous within-host seasonal influenza virus evolution studies (Figure 2A)^26^. Synonymous evolutionary rates were significantly higher than nonsynonymous rates during the initial phase of A/H3N2 virus infections (Figure 2A), primarily in the polymerase complex and HA genes (Figure 2A and S1-2). Importantly, nonsynonymous variants gradually accumulated, increasing in rates around four days post-symptom onset to similar levels relative to synonymous rates. Aggregating over all samples, most nonsynonymous variants were found in the nucleoprotein (NP) and neuraminidase (NA) gene segments (nonsynonymous to synonymous variant (NS/S) ratios = 1.69 (NP) and 1.32 (NA) whereas NS/S ratios were ≤ 1 for all other gene segments; Figure S1 and Table S1). While nonsynonymous NA mutations associated with oseltamivir resistance were positively selected for a subset of individuals in response to the antiviral treatment^21^, nonsynonymous changes to NP were likely mediated by protein stability, T-cell immune response and/or host cellular factors (see next section).

**Figure 2.**
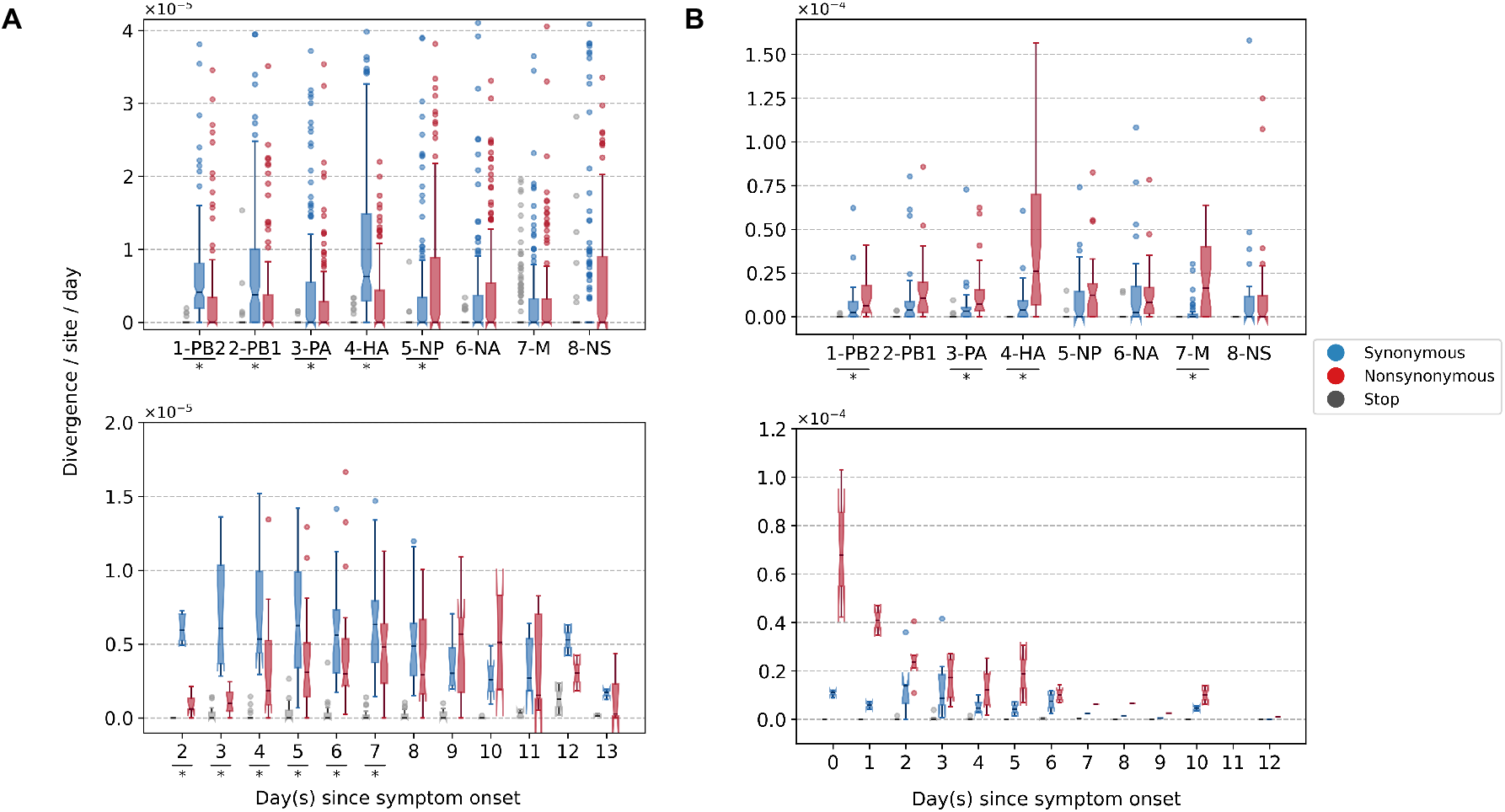
Box plots (median, interquartile range (IQR), and whiskers extending within median ±1.5 × IQR) summarizing the empirical within-host evolutionary rates of (A) seasonal A/H3N2 viruses and (B) pandemic A/H1N1pdm09 viruses. Top panel shows the evolutionary rate of individual gene segments over all timepoints (*r*_*g*_) while the bottom panel depicts the genome-wide evolutionary rate (*r*_*t*_) for each day since symptom onset. All rates are stratified by substitution type (synonymous – blue; nonsynonymous – red; grey – stop-codon). Wilcoxon signed-rank tests were performed to assess if the paired synonymous and nonsynonymous evolutionary rates are significantly distinct per individual gene segment or timepoint (annotated with “ * ” if *p* < 0.05). This was done for all sets of nonsynonymous and synonymous rate pairs except for those computed per day since symptom onset for A/H1N1pdm09 viruses due to the low number of data points available (median number of A/H1N1pdm09 virus samples collected per day since symptom onset = 2). Note that the scales of the y axes differ between A and B to better show rate trends.

For A/H1N1pdm09 viruses during the first wave of the pandemic, the overall within-host evolutionary rate was as high as ∽10^−4^ divergence per site per day in some samples on day 0 post-symptom onset (Figure 2B). Nonsynonymous evolutionary rates were higher than synonymous rates from the start of symptom onset when overall evolutionary rates were also the highest. However, we were unable to determine if the per-day post-symptom onset nonsynonymous and synonymous rates were significantly different from each other due to the low number of samples (i.e. median = 2 samples per day post-symptom onset). Nonetheless, consolidating over all samples across all time points, the polymerase basic 2 (PB2), polymerase acidic (PA), HA and matrix (M) gene segments were the main contributors to the observed rate disparity (Figure 2B and S3-4) with nonsynonymous variants emerging at significantly higher rates relative to synonymous ones. All gene segments also yielded NS/S ratios > 1 (Table S1).

### Intra-host minority variants

Most of the iSNVs identified for both virus subtypes were observed at low frequencies (2-5%; Figure 3), and appear to be stochastically introduced across the virus genome (Figure 4). Purifying selection dominated within-host seasonal A/H3N2 virus populations as the ratio of nonsynonymous to synonymous variants was 0.72 across all samples and variant frequencies (Figures 3A and S2). Of note, the canonical antigenic sites of the HA gene segment^27^ of the A/H3N2 virus populations experienced strong negative selection as evidenced by the occurrence of synonymous variants (median frequency = 0.14, IQR range = 0.09-0.27) at far greater frequencies relative to those at non-antigenic sites of HA (median frequency = 0.03, IQR range = 0.03-0.05; Mann-Whitney U test *p* = 1.18 × 10^−24^; Figure 4C). There were no significant differences in the frequencies of nonsynonymous iSNVs between the antigenic sites of H3 (median frequency = 0.04, IQR range = 0.03-0.06) and the rest of the HA gene segment (median frequency = 0.03, IQR range = 0.02-0.06; Mann-Whitney U test *p* = 0.29; Figure 4C). In contrast, there was 1.94 times as many nonsynonymous minority iSNVs relative to synonymous ones identified in the pandemic A/H1N1pdm09 virus samples (Figures 3B and S4). Variant frequencies of nonsynonymous iSNVs found in the antigenic epitopes of H1^28^ (median frequency = 0.04, IQR range = 0.04-0.05) were, however, not significantly different from those of non-antigenic sites (median frequency = 0.05, IQR range = 0.03-0.16; Mann-Whitney U test *p* = 0.34; Figure 4D).

**Figure 3.**
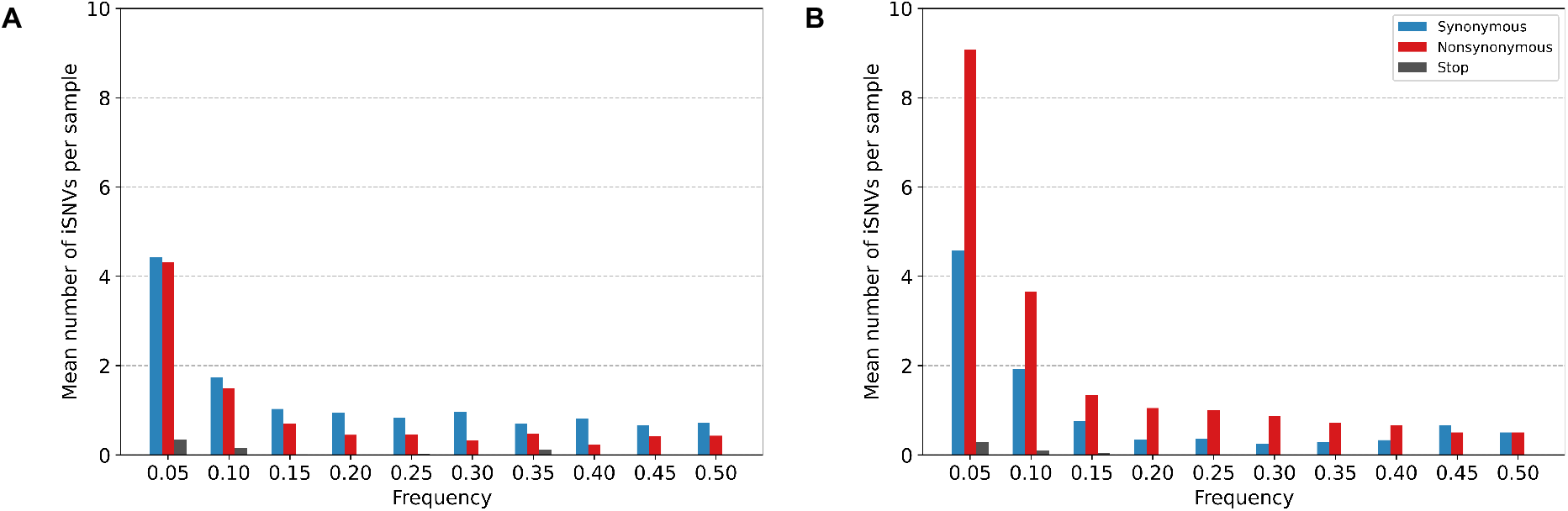
Histogram of the mean number of minority iSNVs identified per sample across all (**A**) A/H3N2 and (**B**) A/H1N1pdm09 virus specimens, sorted by frequency bins of 5% and substitution type (synonymous – blue; nonsynonymous – red; stop-codon – grey).

**Figure 4.**
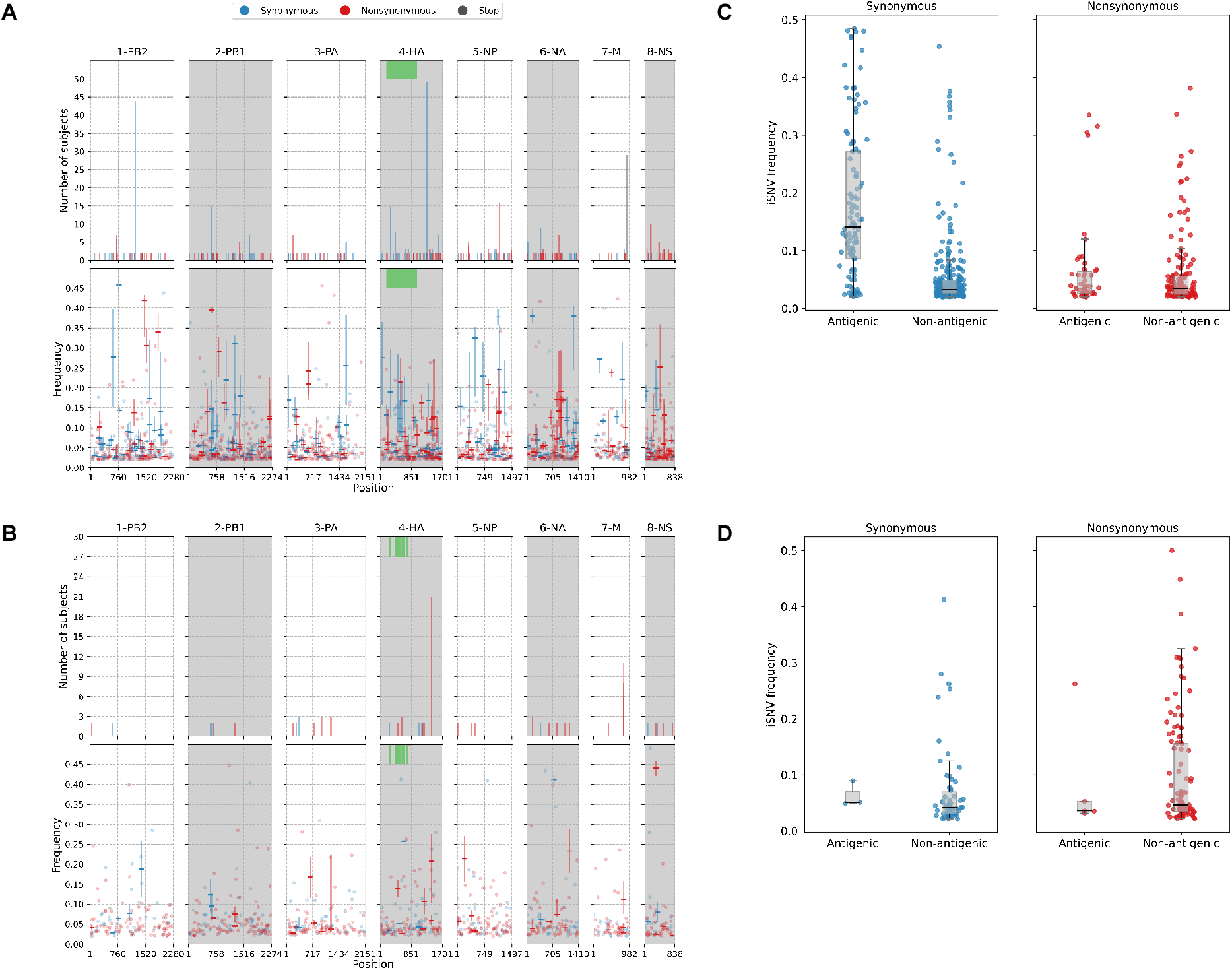
(**A**) Breakdown of iSNVs identified in seasonal A/H3N2 virus samples. The top panels plot the nucleotide positions where iSNVs were found in at least two subjects. The bottom panels shows the frequencies at which iSNVs were identified. For sites with iSNVs that were found in two or more subjects, the interquartile ranges of variant frequencies are plotted as vertical lines and the median frequencies are marked with a dash. If the iSNV was only found in one subject, its corresponding frequency is plotted as a circle. All iSNVs are stratified to either synonymous (blue), nonsynonymous (red) or stop-codon (grey) variants. Only the nonsynonymous variants are plotted if both types of variants are found in a site. Positions of antigenic sites of the haemagglutinin gene segment^29^ are marked in green on the top panels. (**B**) Similar plots to (A) for iSNVs found in pandemic A/H1N1pdm09 virus samples. (**C**) Box plots of the frequencies of synonymous and nonsynonymous variants between antigenic and non-antigenic sites of seasonal A/H3N2 haemagglutinin gene segment. (**D**) Similar plots to (C) for HA iSNVs identified in the pandemic A/H1N1pdm09 virus samples.

As observed in a previous study using different data^26^, premature stop-codon (nonsense) mutations accumulated within-host, though only at low rates. Here, we observed similarly low median nonsense rates, ranging between 0 and 1.2 × 10^−6^ divergence per site per day across the entire A/H3N2 virus genome over the course of infection (IQR limits range between 0 and at most, 1.82 × 10^−6^ divergence per site per day; Figure 2A). Premature stop-codons accumulated in the matrix (M) genes predominantly but also appeared in all other influenza gene segments within various individuals (Figures 2A and 4A). Nonsense mutations also accumulated within the A/H1N1pdm09 virus samples (Figure 2B). Similar to A/H3N2 viruses, nonsense mutation rates were much lower compared to the synonymous and nonsynonymous counterparts (median genome-wide rate across all samples between 0 and 1.43 × 10^−6^ divergence per site per day; IQR limits between 0 and 2.18 × 10^−6^ divergence per site per day).

The premature stop-codon mutations were mostly found at low frequencies for both influenza subtypes (<10%; Figure 3). The exception lies with one of the A/H3N2 virus samples where a premature stop codon was found in position 77 of the M2 ion channel with variant frequency as high as 34.6% (Patient 1843, day 6 since symptom onset; Figures 3A and S5D). The premature stop codon in M2-77 was also found in 27 other individuals across multiple timepoints, albeit at a much lower frequency that never amounted more than 10% (Figures 4A and S5D). This was unlikely to be a sequencing artefact resulting from a mistaken incorporation of the primer sequence as its carboxyl terminal falls outside the coding region of the M gene segment (Table S3) and the variant frequencies would have been much higher in all samples if this was the case.

Despite the dominance of purifying selection in seasonal A/H3N2 intra-host viral populations, we detected several nonsynonymous variants of interest. Amino acid variants emerging in the HA and NA proteins were discussed in a previous work^21^ (see Supplementary Materials). In the nucleoprotein, there were two notable nonsynonymous variants, D101N/G and G384R, that appeared in multiple individuals who were sampled independently between 2007 and 2009 (Figure 4A and S5C). D101N/G was found in 7 different patients and at least for D101G, the mutation was previously linked to facilitating escape from MxA, a key human antiviral protein^30^. However, the nonsynonymous mutation was only found in low frequencies and remained invariant during the respective courses of infection for all seven patients (median variant frequency across all samples = 0.03; IQR = 0.02-0.07).

NP-G384R emerged in sixteen unlinked patients infected by A/H3N2 virus. Even though G384R did not become the majority variant in any of these individuals (median variant frequency across all samples = 0.14; IQR = 0.07-0.20), the variant emerged around day 4-5 post-symptom onset and mostly persisted within each individual for the rest of sampled timepoints. G384R is a stabilizing mutation in the A/Brisbane/10/2007 A/H3N2 virus NP background^31^ that is similar to the viruses investigated here. Interestingly, position 384 is an anchor residue for several NP-specific epitopes recognised by specific cytotoxic T lymphocytes (CTLs) that are under continual selective pressure for CTL escape^32,33^. The wild-type glycine residue is known to be highly deleterious even though it was shown to confer CTL escape among HLA-B *2705-positive individuals^34–36^.

Using a maximum likelihood approach to reconstruct and estimate the frequencies of the most parsimonious haplotypes of each gene segment, we computed linkage disequilibrium and found evidence of potential epistatic co-variants to NP-G384R in the A/H3N2 virus populations of multiple individuals (Figure 5 and Table S2). When analysing how these variants could alter protein stability using FoldX, the stabilizing effects of G384R (mean Δ Δ = −3.84 kcal/mol (SD = 0.06 kcal/mol)) was found to alleviate the likely destabilizing phenotype of a functionally relevant linked variant in two of the three co-mutation pairs identified in separate individuals (i.e. G384R/M426I and G384R/G102R; Table S2). In the first individual (subject 1224), M426I was inferred to have emerged among the viral haplotypes encoding NP-G384R on the 10^th^ day post-symptom onset (D10). M426I may be compensating for T-cell escape that was previously conferred by 384G even though the two amino acid sites are anchor residues of different NP-specific CTL epitopes^32^. M426I was found to be highly destabilizing (mean Δ Δ = 2.61 kcal/mol (standard deviation (SD) = 0.05 kcal/mol); Table 1) but when co-mutated with G384R, stability changes to NP was predicted to be neutral (mean Δ Δ = − 0.42 kcal/mol (SD = 0.06 kcal/mol)). In the second individual (subject 1686), G102R was likely linked to G384R in the within-host virus populations found in the D10 sample. As a single mutant, G102R is also destabilizing to NP (mean Δ Δ = 4.87 kcal/mol (SD = 0.00 kcal/mol)). However, when combined with G384R, NP protein stability was only weakly destabilizing (mean Δ Δ = 0. 6 kcal/mol (SD = 0.09 kcal/mol)). G102R was previously found to bypass the need for cellular factor importin-α7 which is crucial for viral replication and pathogenicity of IAVs in humans^37–39^.

**Table 1:** FoldX stability predictions of likely linked nonsynonymous minority variants found in A/H3N2 nucleoprotein. The mean ΔΔ*G* and standard deviation (S.D.) values reported are based on the results of five distinct simulations. Variants with mean ΔΔ*G* < − 0.46 kcal/mol are deemed to be stabilizing while destabilizing mutants were estimated to yield ΔΔ*G* > 0.46 kcal/mol.

| Variants | $\Delta\Delta G$ (kcal/mol) | |
| --- | --- | --- |
|  | Mean | S.D. |
| G384R | -3.84 | 0.06 |
| M426I | 2.61 | 0.05 |
| G384R,M426I | -0.42 | 0.06 |
| G102R | 4.87 | 0.00 |
| G384R,G102R | 0.76 | 0.09 |
| A493T | 11.96 | 0.30 |
| G384R,A493T | 5.56 | 0.19 |
| V197I | -3.11 | 0.02 |
| S353Y | -1.97 | 0.68 |
| V197I,S353Y | -4.48 | 0.14 |

**Figure 5.**
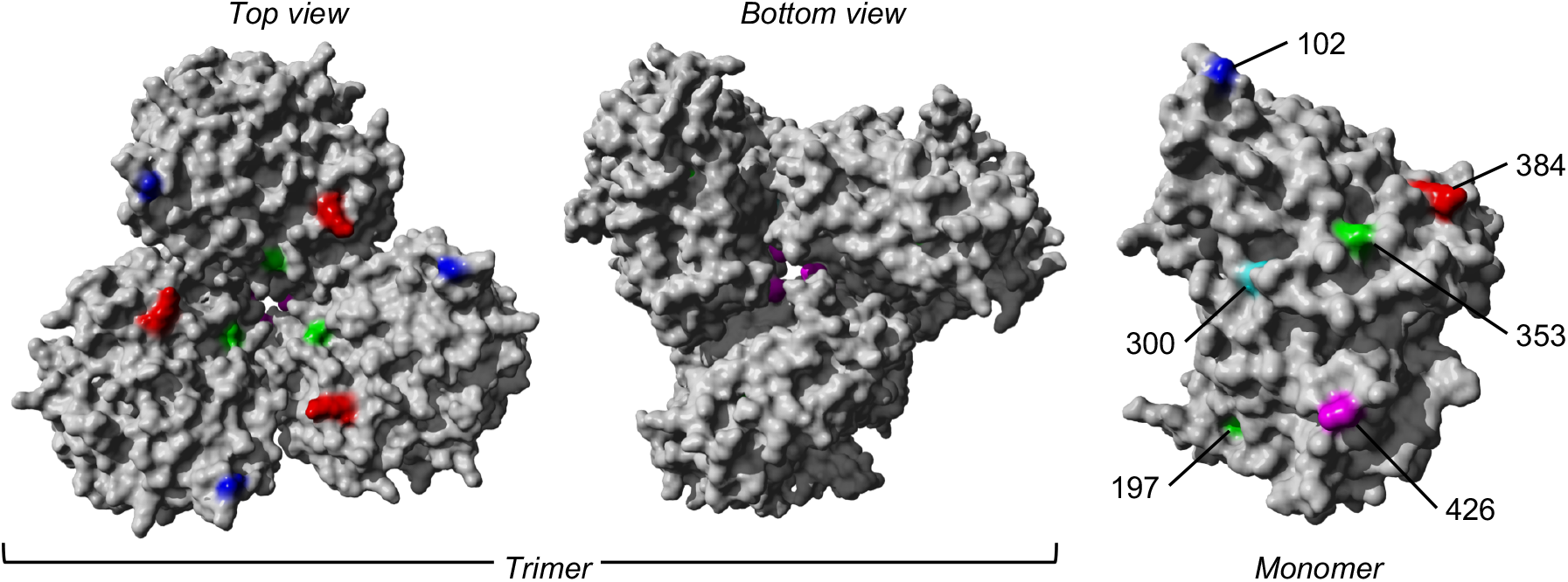
The trimeric and monomeric crystal structures of nucleoprotein (PDB: 3ZDP)^40^ of influenza A viruses. Amino acid sites with potentially linked epistatic amino acid variants as tabulated in Table 1 are separately coloured, with their corresponding positions annotated on the monomeric structure.

For the pandemic A/H1N1pdm09 viruses, most of the nonsynonymous variants were found singularly in individual patients (Figure 4B). Putative HA antigenic minority variants were found in four individuals in distinct amino acid sites (G143E, N159K, N197K and G225D; H3 numbering without signal peptide; Figure S5E). All of these variants were found at frequencies ≤5% and the wild-type residues have been conserved in the corresponding positions globally to date, with the exception of position 225. Here, HA-225G was the majority variant (76%) in a hospitalised individual (subject 11-1022; Table S4) and D225G is linked to infections with severe disease outcomes^41^. Furthermore, one of the few nonsynonymous iSNVs that co-emerged in multiple unlinked patients was found in the usually conserved stem of the HA protein, L455F/I (H3 numbering without signal peptide), appearing in 17 separate individuals (Figures 4B and S5E). The amino acid variant was found in patients from different time periods and geographical locations (Table S4), thus it is unlikely this was a unique variant shared among individuals in the same transmission cluster. It was observed as early as day 0 post-symptom onset for some patients and seemed to persist during the infection but only as a minority variant at varying frequencies (median frequency across all samples with mutation = 0.20; IQR = 0.08-0.28). However, this position has also been conserved with the wild-type Leucine residue in the global virus population to date. Hence, it is unclear if HA-L455F/I actually confers any selective benefit even though it was independently found in multiple patients.

We also found oseltamivir resistance mutation H275Y^42^ in the NA proteins in two unlinked individuals who were infected with the A/H1N1pdm09 virus and treated with oseltamivir (Figure S5F and Table S4). 275Y quickly became the majority variant in both patients within 3-4 days after the antiviral drug was first administered. Finally, there were two other amino acid variants in the M2 ion channel that appeared within multiple subjects in parallel across different geographical locations – L46P and F48S were identified in 8 and 16 patients respectively in a range of frequencies (L46P: median frequency = 0.04, IQR = 0.04-0.05; F48S: median frequency = 0.08, IQR = 0.03-0.13) but similarly, never becoming a majority variant in any of them (Figures 4B and S5G). Again, the wild-type residues were mostly conserved in the global virus population since the pandemic.

### Within-host simulations

To investigate the evolutionary pressures that likely underpin the observed patterns of synonymous and nonsynonymous substitutions (Figure 2), we performed forward-time Monte Carlo simulations. Given that the median age of the children infected by A/H3N2 virus at the time of sample collection was 2 years of age (IQR=2-3 years), most of them were likely experiencing one of their first influenza virus infections. Furthermore, influenza vaccination for children is not part of the national vaccination programme in Vietnam. As such, most of the children analysed here lacked influenza virus specific antibodies based on haemagglutination inhibition assays^21^. For individuals infected by the pandemic A/H1N1pdm09 virus, all but one patient was under 60 years of age and thus lacked immunity to the virus as well. Furthermore, patients infected by either viruses mount little-to-no humoral immune selection pressure during the first 7-10 days of infection^43^. As such, the contrasting evolutionary dynamics between these viruses (Figure 2) are unlikely to have been driven by antibody-mediated selection pressure.

Seasonal A/H3N2 viruses, having circulated within the human population since 1968, are expected to be well adapted to human hosts at this point such that most nonsynonymous mutations are likely highly deleterious and would not reach detectable frequencies. Those that were detected are mostly expected to be weakly deleterious, and thus not purged fast enough by selection such that mutation-selection balance was observed. In contrast, there was evolutionary space for A/H1N1pdm09 virus to further adapt to its new found human hosts during the initial waves of the pandemic. Since no mutation selected by rapid directed positive selection was observed, most of the detected nonsynonymous mutations were expected to be neutral and a small but non-trivial fraction are likely to be weakly beneficial.

Our simulations used a simple within-host evolution model represented by a binary genome that distinguishes between synonymous and nonsynonymous loci. Given that the estimated transmission bottleneck sizes for pandemic A/H1N1pdm09 (see Supplementary Materials) and seasonal A/H3N2 viruses^4,44^ are narrow at 1-2 genomes, we modelled an expanding virus population size during the initial timepoints of the infection that started with one virion. If within-host virus populations were to evolve neutrally, we would observe similar synonymous and nonsynonymous evolutionary rates throughout the infection (Figure 6C). On the other hand, if selection is sufficiently strong, accumulation of beneficial (or deleterious) nonsynonymous variants will increase (or decrease) substantially with time (Figure 6B and 6E). Clearly, these patterns were not observed for both IAVs (Figure 2).

**Figure 6:**
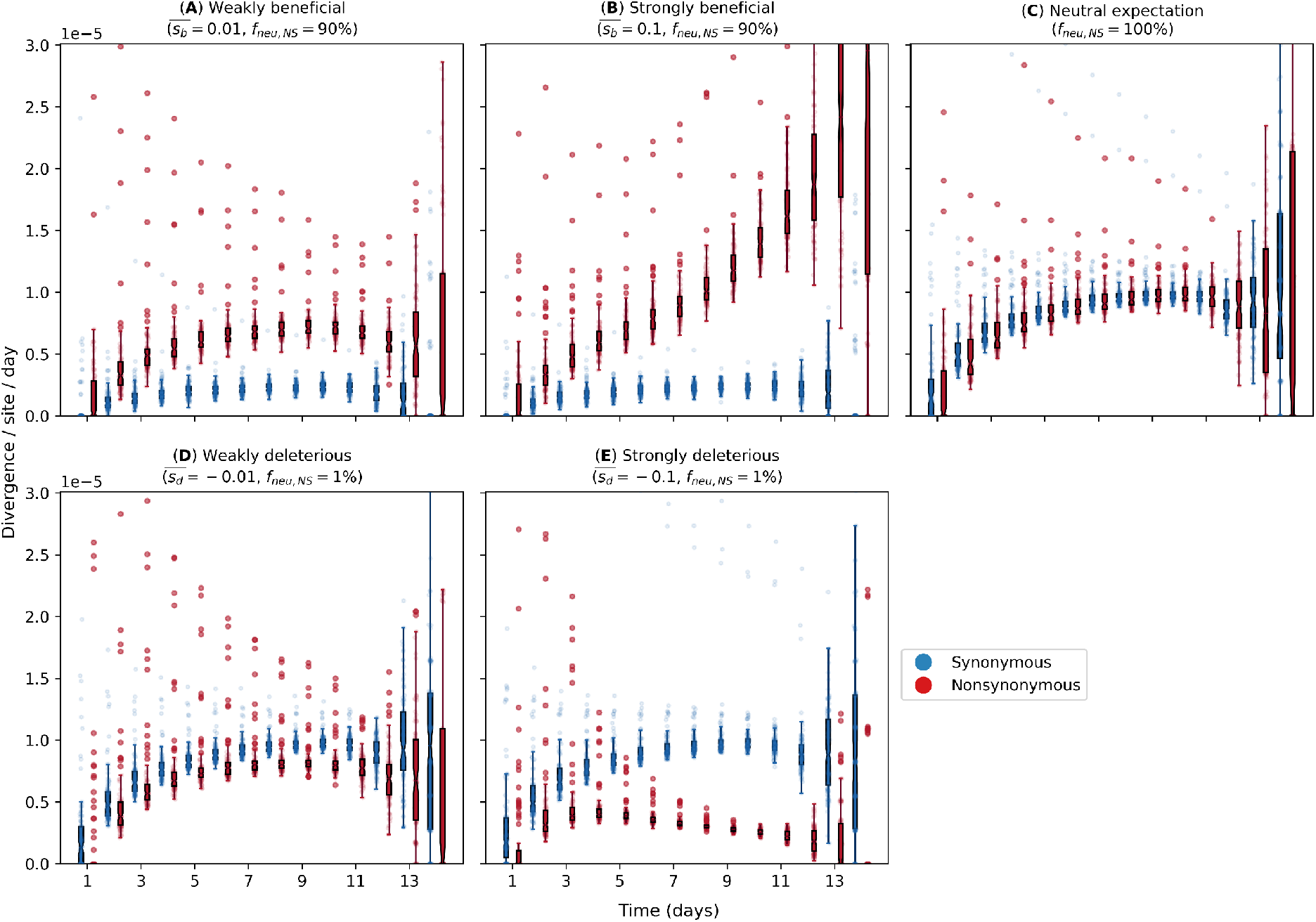
Evolutionary rates computed from forward-time Monte Carlo within-host simulations for different fitness effects of nonsynonymous mutations (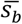 and 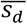 denote mean beneficial and deleterious effects respectively) and fraction of neutral nonsynonymous mutations (*f* _*neu,NS*_). We assumed that synonymous mutations are neutral for all simulations. For A/H1N1pdm09 viruses, we assumed that only a small fraction of nonsynonymous mutations is neutral (*f* _*neu,NS*_ = 1%) and performed simulations where the remaining nonsynonymous mutations are either (**A**) weakly 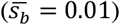 or (**B**) strongly 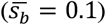 beneficial. For A/H3N2 viruses, we tested the hypotheses where majority of nonsynonymous mutations are neutral (*f* _*neu,NS*_ = 0%) while the remaining ones are either (**D**) weakly 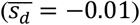 or (**E**) strongly 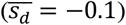 deleterious. (**C**) Neutral expectation where all nonsynonymous mutations are neutral (*f* _*neu,NS*_= 100%).

However, if most *de novo* nonsynonymous mutations are only weakly deleterious, we would observe larger synonymous evolutionary rates initially before nonsynonymous variants accumulate to similar levels (Figure 6A). By then, virion population size (*N*) would also be large enough relative to the virus mutation rate (μ) (i.e.*N*μ ≫1; see Supplementary Materials) such that mutation-selection balance is expected and evolutionary rates remain fairly constant, similar to the patterns empirically observed for within-host A/H3N2 virus populations (Figure 2A). Contrastingly, if the majority of nonsynonymous variants are neutral and only a small subset confers weakly beneficial effects, nonsynonymous evolutionary rates would consistently be larger than their synonymous counterpart but never accumulate to levels akin to those observed for strong positive selection (Figure 6D). Although the simulation results here does not entirely reflect the evolutionary dynamics observed for A/H1N1pdm09 viruses in Figure 2B, we hypothesised that there was substantial virus replication prior to symptom onset and that our samples better reflect the virus populations present midway or nearing the end of the infection when compared to our simulation results. This is further evidenced by the relatively large number of iSNVs detected at the time of symptom onset (Figure 1B) and the tight transmission bottleneck sizes we estimated for the pandemic virus (see Supplementary Materials).

## Discussion

Multiple next-generation sequencing studies have found little evidence of positive selection in seasonal influenza virus populations of acutely infected individuals^4,8–11,45^. Recent modelling work showed that the time required to initiate new antibody production and asynchrony with virus exponential growth limits the selection of *de novo* antigenic variants within host in acute seasonal influenza virus infections^46^. In contrast, phenotypically relevant variants that were positively selected in within-host virus populations of severely immunocompromised patients coincided with those selected by the global seasonal IAV population^47,48^. This implies that within-host evolutionary dynamics of seasonal IAVs in immunocompromised individuals are likely to be substantially different owing to the increased time for virus diversity to accumulate and for selection to act^49^. In other words, the duration of infection is likely to be critical for positive evolutionary selection to be effective within host.

Viral shedding duration is often longer in young children infected with seasonal influenza virus compared to otherwise healthy adults^50^. Children also play a critical role in “driving” influenza epidemics due to their higher contact and transmission rates^22,23^. As such, our seasonal A/H3N2 virus results fill an important gap in the current literature of within-host evolutionary studies of seasonal IAVs as most of the samples analysed were collected from children under the age of six years up to two weeks post-symptom onset. Importantly, the absence of antibody-mediated immunity in young unvaccinated children, which would otherwise reduce the extended duration of infection, has the potential to facilitate other routes of virus evolution.

Similar to the aforementioned within-host studies, the A/H3N2 virus population within these children was characterised by low genetic diversity and dominated by purifying selection early in the infection. Due to a lack of antibody response against the antigenic regions of HA^21^, it is unsurprising that we observed a lack of adaptive changes to the HA antigenic regions, similar to adults in previous studies^4^. We also found that the polymerase genes were subjected to purifying selection, indicating their critical role in virus replication as negative selection purges deleterious variation. However, while purifying selection is detectable, it is incomplete^26^. We observed that most nonsynonymous variants began to accumulate around 3-4 days post-symptom onset, with incrementally higher empirical rates as the infection progressed.

Through simulations of a within-host evolution model, we hypothesised that the accumulation of nonsynonymous iSNVs was a result of their weakly deleterious effects and expanding virion population size such that mutation-selection balance was reached. The maintenance of genetic diversity through mutation-selection balance within these children may provide opportunities for the emergence of phenotypically relevant mutations which deleterious effects could be alleviated by the accumulation of a secondary compensatory mutations. For example, in one individual NP-G384R was accompanied by NP-M426I which is an anchor residue of a CTL epitope of NP, abrogating recognition by HLA-B *3501-positive CTLs^32^ but is likely to be deleterious based on our computational protein stability predictions. G384R, which is located in a CTL epitope distinct from M426I^32^, was previously shown to be a stabilizing substitution^31^.

Interestingly, we also observed G384R in the minority virus population of 15 other unlinked individuals. Besides improving NP protein stability, G384R restores recognition by HLA-B *2705-positive NP-specific CTLs^36^. The NP gene segment in the global A/H3N2 virus population has an evolutionary history of fixating destabilizing amino acid mutations that promote CTL immune escape alongside stabilizing substitutions that compensate for the deleterious effects of the former^34^. The reversal R384G mutation confers CTL escape but is known to be highly deleterious. This substitution was fixed in the global A/H3N2 virus population during the early 1990s as other substitutions such as S259L and E375G epistatically alleviated its destabilizing effects^34^. One possible explanation for the emergence of G384R as a minority variant within these unlinked individuals is that they are all HLA-B *2705 negative. However, we did not collect the necessary blood samples to investigate this possibility.

In contrast, we found a substantially higher fraction of nonsynonymous variants in the within-host virus populations of individuals infected A/H1N1pdm09 virus during the pandemic. Owing to the different next-generation sequencing platforms used to sequence samples of the two virus subtypes and consequently differences in base calling error rates and depth of coverage (Figure S6), we did not directly compare the observed levels of within-host genetic diversity between the two influenza subtypes here. However, given that only iSNVs with frequencies ≥ 2% were called, low-frequency minority variants arising from technical-related errors should be minimised^51^. Importantly, the relative number of nonsynonymous iSNVs identified were far greater than synonymous ones early in the pandemic A/H1N1pdm09 virus infections, suggesting that there was room for further human host adaptation, particularly in the HA but also in the polymerase gene segments similar to those observed in other zoonotic influenza virus infections^52^.

Given the tight estimated transmission bottleneck size (see Supplementary Materials), the relatively large number of iSNVs identified at the start of symptom onset and simulations of within-host evolution (see Supplementary Materials, Figure 1B, 2B and 6D), it is unlikely that the initial within-host A/H1N1pdm09 virus populations sampled were the inoculating population that founded the infection. Instead, the inoculating viral population had already undergone substantial within-host replication during the incubation period before symptom-onset. In fact, four of the individuals analysed were asymptomatic (i.e. H058/S02, H089/S04, H186/S05 and H296/S04; Table S4). Additionally, pre-symptomatic virus shedding was observed in some of the secondary household cases^53^ and presymptomatic transmission has been documented in other settings^54^. Nonetheless, this would not meaningfully impact our conclusions as most of the within-host viral populations sampled at the start of symptom onset should still constitute those found early in infection and the contrasting feature where nonsynonymous iSNVs outnumbered synonymous ones were not observed in the seasonal A/H3N2 virus samples.

For both A/H3N2 and A/H1N1pdm09 virus samples, nonsense iSNVs resulting in premature stop codons were found to accumulate within host, even though only at low proportions. The accumulation of premature stop-codon mutations further suggest that while purifying selection dominates within-host influenza virus populations, it may not be acting strongly enough to completely purge these lethal nonsense mutations^26^. Additionally, it has been recently found that incomplete influenza virus genomes frequently occur at the cellular level and that efficient infection depends on the complementation between different incomplete genomes^55^. As such, nonsense mutations may not be as uncommon as previously thought. In particular, nonsense mutations in position 77 of the M2 ion channel were independently found in 27 unlinked individuals infected by A/H3N2 virus. While these nonsense mutations are generally considered to be lethal, ion channel activity is retained even if the M2 protein was prematurely truncated up to position 70 at its cytoplasmic tail^56^.

Our study has several limitations. The number of iSNVs identified can potentially be biased by variations in sequencing coverage^57^. As such, the number of iSNVs observed in one intra-host virus populations may not be directly comparable to another with a distinct coverage profile (Figure S6). As an alternative, the nucleotide diversity π statistic^58^ may be a more robust measure of within-host diversity as it solely depends on the underlying variant frequencies^57^. Computing the corresponding π statistics for our data, we observed trends in genetic diversity that were similar to those inferred using iSNV counts (see Supplementary Materials and Figure S7).

To ensure accurate measurements of virus diversity in intra-host populations, we would also need to be certain that the estimated variant frequencies precisely reflect the distributions of variants that comprise the sampled virus populations. The inferred variant frequencies can be significantly distorted if virus load is low^59,60^. As such, we limited our analyses for both virus subtypes to samples with Ct-values ≤35 which likely afford sufficient virus material for sequencing^60^. We were unable to estimate the amount of frequency estimation errors for the A/H1N1pdm09 virus samples as only one sequencing replicate was performed using the universal 8-segment PCR method^61^. However, for the A/H3N2 virus samples, independent PCR reactions were performed using three partly overlapping amplicons for all gene segments other than the non-structural and matrix genes. We compared the variant frequencies estimated for any overlapping sites generated by reads derived from distinct amplicons with sufficient coverage (>100x). Variant frequencies computed from independent amplicons agreed well with each other across the range of Ct values of the samples from which variants were identified (Figure S8), affirming the precision of our iSNV frequency estimates for the A/H3N2 virus samples, including those with higher Ct values.

Finally, most study participants received oseltamivir during the course of their infections (Table S4). Although we were unable to identify any potential effects of enhanced viral clearance or any other evolutionary effects due to the treatment, besides oseltamivir-resistance associated mutations, it is unlikely that the antiviral treatment had a substantial impact on our results. First, the median timepoint in which the antiviral was initially administered was 4 days post-symptom onset (IQR = 3-6 days; Table S4). Previous studies showed that enhanced viral clearance of IAVs was mostly observed among patients who were treated with oseltamivir within 3 days of symptom onset^20,62,63^. Of note, late timepoint samples in this study (≥ 8 days since symptom onset) mostly came from individuals who started oseltamivir treatments ≥ 4 days post-symptom onset (Figure S12). Second, at least *in vitro*, there were no differences in the levels of genetic diversity observed in influenza virus populations after multiple serial passages whether they were treated with oseltamivir or not^64^.

To conclude, we presented how intra-host populations of seasonal and pandemic influenza viruses are subjected to contrasting evolutionary selection pressures. In particular, we showed that the evolutionary dynamics and ensuing genetic variation of these within-host virus populations changes during the course of infection, highlighting the importance for sequential sampling, particularly for longer-than-average infections such as those in the young children studied here.

## Methods

### Sample collection and viral sequencing

The A/H3N2 virus samples were collected from 52 patients between August 2007 and September 2009 as part of an oseltamivir dosage trial conducted by the South East Asia Infectious Disease Clinical Research Network (SEAICRN), which is detailed in a previous work^20^. Briefly, patients with laboratory confirmed influenza virus infection and duration of symptoms ≤ 10 days were swabbed for nose and throat samples daily between 0 and 10 days as well as day 14 upon enrolment for the study (Table S4). All PCR-confirmed A/H3N2 virus samples with cycle threshold (Ct) values ≤ 35 were included for sequencing.

Library preparation and viral sequencing protocols performed on these A/H3N2 virus samples are elaborated in detail in ^21^. Here, we highlight key aspects of our preparation and sequencing procedures. Using segment specific primers (Table S3), we performed six independent PCR reactions, resulting in three partly-overlapping amplicons for each influenza virus gene segment other than the matrix (M) and non-structural (NS) genes where a single amplicon was produced to cover the entirety of the relatively shorter M and NS genes. The use of shorter but overlapping amplicons in the longer gene segments improve amplification efficiency, ensuring that these longer segments are sufficiently covered should there be any RNA degradation in the clinical specimen. These overlapping PCR products were pooled in equimolar concentrations for each sample and purified for subsequent library preparation. Sequencing libraries were prepared using the Nextera XT DNA Library Preparation kit (Illumina, FC-131-1096) as described in ^21^. Library pools were sequenced using the Illumina MiSeq 600-cycle MiSeq Reagent Kit v3 (Illumina, MS-102-3003).

The A/H1N1pdm09 virus samples were obtained as part of a household-based influenza virus cohort study that was also performed by SEAICRN. The study was conducted between July and December 2009, involving a total of 270 households in Ha Nam province, Vietnam^24^. Similarly, combined nose and throat swabs were collected daily for 10-15 days from individuals with influenza-like-illness (i.e. presenting symptoms of fever >38°C and cough, or sore throat) and their household members, including asymptomatic individuals (Table S4). We also analysed additional samples collected from unlinked hospitalised patients who were infected by the A/H1N1pdm09 virus from two major Vietnamese cities (Hanoi and Ho Chih Minh) during the first wave of the pandemic^20,25^. A total of 32 PCR-confirmed A/H1N1pdm09-infected individuals originating from both households and hospitalised cases were selected for sequencing based on availability and Ct-values ≤ 33 (Table S4).

For the A/H1N1pdm09 virus samples, RNA extraction was performed manually using the High Pure RNA isolation kit (Roche) with an on-column DNase treatment according to the manufacturer’s protocol. Total RNA was eluted in a volume of 50 μl. Universal influenza virus full-genome amplification was performed using a universal 8-segment PCR method as described previously^51,65,66^. In short, two separate RT-PCRs were performed for each sample, using primers common-uni12R (5’-GCCGGAGCTCTGCAGAT ATCAGCRAAAGCAGG-3’), common-uni12G (5’-GCCGGAGCTCTG CAGATATCAGCGAAAGCAGG-3’), and common-uni13 (5’-CAGGAA ACAGCTATGACAGTAGAAACAAGG-3’). The first RT-PCR mixture contained the primers common-uni12R and common-uni13. The second RT-PCR mixture contained the primers common-uni12G and common-uni13, which greatly improved the amplification of the PB2, PB1, and PA segments. Reactions were performed using the One-Step RT-PCR kit High Fidelity (Invitrogen) in a volume of 50 μl containing 5.0 μl eluted RNA with final concentrations of 1xSuperScript III One-Step RT-PCR buffer, 0.2 μM of each primer, and 1.0 μl SuperScript III RT/Platinum Taq High Fidelity Enzyme Mix (Invitrogen). Thermal cycling conditions were as follows: reverse transcription at 42°C for 15 min, 55°C for 15 min, and 60°C for 5 min; initial denaturation/enzyme activation of 94°C for 2 min; 5 cycles of 94°C for 30 s, 45°C for 30 s, slow ramp (0.5°C/s) to 68°C, and 68°C for 3 min; 30 cycles of 94°C for 30 s, 57°C for 30 s, and 68°C for 3 min; and a final extension of 68°C for 5 min. After the PCR, equal volumes of the two reaction mixtures were combined to produce a well-distributed mixture of all 8 influenza virus segments. All RT-PCRs were performed in duplicate. Samples were diluted to a DNA concentration of 50 ng/µl followed by ligation of 454 sequencing adaptors and molecular identifier (MID) tags using the SPRIworks Fragment Library System II for Roche GS FLX+ DNA Sequencer (Beckman Coulter), excluding fragments smaller than 350 base pairs, according to the manufacturers protocol to allow for multiplex sequencing per region. The quantity of properly ligated fragments was determined based on the incorporation efficiency of the fluorescent primers using FLUOstar OPTIMA (BMG Labtech). Emulsion PCR, bead recovery and enrichment were performed manually according to the manufacturers protocol (Roche) and samples were sequenced in Roche FLX+ 454. Sequencing was performed at the Sanger Institute, Hinxton, Cambridge, England as part of the FP7 program EMPERIE. Standard flowgram format (sff) files containing the filter passed reads were demultiplexed based on the molecular identifier (MID) sequences using QUASR package version 7.0^51^.

### Read mapping

Trimmomatic (v0.39; Bolger et al. 2014) was used to discard reads with length <30 bases while trimming the ends of reads where base quality scores fall below 20. The MAXINFO option was used to perform adaptive quality trimming, balancing the trade-off between longer read length and tolerance of base calling errors (target length=40, strictness=0.4). For the A/H3N2 virus samples, the trimmed paired reads were merged using FLASH (v1.2.11)^68^. All remaining reads were then locally aligned to A/Brisbane/10/2007 genome (GISAID accession: EPI_ISL_103644) for A/H3N2 virus samples and A/California/4/2009 genome (EPI_ISL_376192) for A/H1N1pdm09 virus samples using Bowtie2 (v2.3.5.1)^69^. Aligned reads with mapping scores falling below 20 alongside bases with quality score (*Q-score*) below 20 were discarded.

### Variant calling and quality filters

Minority variants of each nucleotide site with a frequency of at least 2% were called if the nucleotide position was covered at least 50x (H1N1pdm09) or 100x (H3N2) and the probability that the variant was called as a result of base calling errors (*p*_*Err*_) was less than 1%. *p*_*Err*_ was modelled by binomial trials^70^:

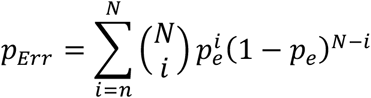

where 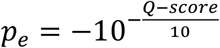, *N* is the coverage of the nucleotide site in question and *n* is the absolute count of the variant base tallied.

While lower coverage at both ends of individual gene segments was expected, there were also variable coverage results across gene segments for some samples that were mapped to A/H3N2 virus (Figure S6). In order to retain as many samples deemed to have adequate coverage across whole genome, a list of polymorphic nucleotide sites found to have >2% minority variants in more than 1 sample was compiled. Each gene segment of a sample was determined to achieve satisfactory coverage if >70% of these polymorphic sites were covered at least 100x. For A/H1N1pdm09, the gene segment of a sample was deemed to be adequately covered if 80% of the gene was covered at least 50x.

The number of iSNVs observed in A/H3N2 virus samples collected from subject 1673 (39-94 iSNVs in three samples collected from three (D3) to five (D5) days post-symptom onset) and the D8 sample for subject 1878 (73 iSNVs) were substantially greater than numbers in all other samples. The putative majority and minority segment-concatenated sequences of these samples did not cluster as a monophyletic clade among themselves phylogenetically (Figure S9), suggesting that these samples might be the product of mixed infections or cross-contamination. These samples were consequently excluded from further analyses.

### Empirical within-host evolutionary rate

The empirical within-host evolutionary rate (*r*_*g,t*_) of each gene segment (*g*) in a sample collected on *t* day(s) since symptom onset were estimated by:

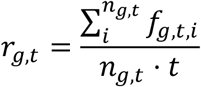

where *f*_*g,t,i*_ is the frequency of minority variants present in nucleotide site *i* for gene segment *g* and *n*_*g,t*_ is the number of all available sites^26^. Distinct rates were calculated for synonymous and non-synonymous iSNVs. The corresponding whole-genome evolutionary rate (*r*_*t*_) on day is computed by summing the rates across all gene segments:

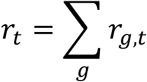

### Within-host simulations

We implemented forward-time Monte Carlo simulations with varying population size using a simplified within-host evolution model to test if our hypotheses could explain the different evolutionary dynamics observed between A/H3N2 and A/H1N1 viral populations. We assumed that a single virion leads to a productive influenza virus infection within an individual and computed changes in the virus population size (*N*) using a target cell-limited model. New virions are produced upon infection by existing virions at a rate of β*CN* where *C* is the existing number of target cells while β is the rate of per-cell per-virion infectious contact. Upon infection, a cell will produce *r* number of virions before it is rendered unproductive. We assume that infected individuals did not mount any antibody-mediated immune response, setting the virus’ natural per-capita decay rate (*d*) such that virions continue to be present within host for 14 days (Figure S11 and Table 2). β is then computed by fixing the within-host basic reproduction number (*R*_0_):

**Table 2:** Parameter values used in within-host model

| Parameter | Meaning | Value (units) | Source |
| --- | --- | --- | --- |
| - | Number of hours per replicative generation | 6 hours | Assumption |
| $r$ | Average number of virions produced by an infected cell | 100 virions | 71 |
| $C_0$ | Initial target cell population size | $4 \times 10^8$ virions | 72 |
| $d$ | Per-capita decay rate | 2 per-generation | Assumption |
| $R_0$ | Within-host basic reproduction number | 5 | 72 |
| $\mu$ | Per-site, per-generation mutation rate | $3 \times 10^{-5}$ per-site, per-generation | 45 |

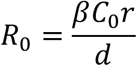

where *C*_0_ is the initial (maximum) target cell population size. We solve the following system of ordinary differential equations numerically to compute the number of virions per viral replicative generation (*N*(*t*)):

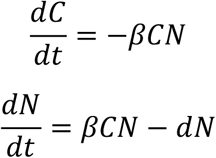

We assume a binary genome of length *L*, distinguishing between synonymous and nonsynonymous loci. For A/H3N2 viruses, we hypothesised that most *de novo* mutations are either weakly deleterious or neutral. To estimate the number of such sites, we aligned A/H3N2 virus sequences that were collected between 2007 and 2012 and identified all polymorphic sites with variants that did not fixate over time (i.e. <95% frequency over one-month intervals). We estimated *L* = 1050 with 838 and 212 synonymous and nonsynonymous loci respectively. On the other hand, for A/H1N1pdm09 viruses, we assumed that any variants that emerged are likely neutral or weakly beneficial. In the absence of strong purifying selection, ∼75% of mutations are expected to be nonsynonymous^26^. Here, we assumed *L* = 1000 sites of which 750 of them are synonymous and the rest are nonsynonymous.

We tracked the frequency distribution of genotypes present for every generation *t*. We assumed that mutations occur at per-locus, per-generation rate μ. During each generation *t*, the number of virions incurring a single-locus mutation followed a Poisson distribution with mean *N* (*t*) μ*L*. For each virion, the mutant locus was randomly selected across all loci. We assumed that all synonymous and a fraction of nonsynonymous sites (*f*_*neu,NS*_) are neutral (i.e. (log) fitness effect = 0). The remaining nonsynonymous sites either had an additive deleterious (*s*_*d*_) or beneficial (*s*_*b*_) fitness effect when mutated. The magnitude of *s_d_* / *s_b_* follow an exponential distribution with mean effect 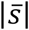. Epistasis was neglected throughout. The distribution of genotypes in the next generation *t* +1 was achieved by resampling individuals according to Poisson distribution with mean *N*(*t* + 1)*p*_*f*_(*g*.*t*) where *p*_*f*_(*g*.*t*) is the relative fitness distribution of genotype *g* during generation *t*.

To decrease the computational costs of the simulations, specifically when *N*(*t*) reaches orders of 10^10^ − 10^11^ virions (Figure S11), we implemented an upper population size limit of 10^7^ virions. Given the mutation rate assumed (Table 2), *N*(*t*) μ ≫1 for *N*(*t*) ≥ 10^7^ virions, mutation-selection balance is theoretically expected for a single-locus (deleterious) mutant model (see Supplementary Materials). We ran 500 simulations for each variable set of *f*_*neu,NS*_and *s_d_* / *s_b_* values. All parameter values used in the model are given in Table 2.

### Haplotype reconstruction

The most parsimonious viral haplotypes of each gene segment were reconstructed by fitting the observed nucleotide variant count data to a Dirichlet multinomial model using a previously developed maximum likelihood approach to infer haplotype frequencies^44^. Assuming that the viral population is made up of a set of *K* haplotypes with frequencies ***q***_***k***_, the observed partial haplotype frequencies ***q***_***l***_ for a polymorphic site can be computed by multiplying a projection matrix ***T***_***k***_. For instance, if the set of hypothetical full haplotypes is assumed to be {*AA,GA,AG*}, the observed partial haplotype frequencies for site *l* = 1, *q* _*A−*_ and *q* _*G*−_ are computed as:

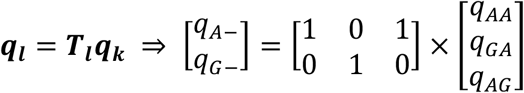

A list of potential full haplotypes was generated from all combinations of nucleotide variants observed in all polymorphic sites of the gene segment. Starting from *K* = 1 full haplotype, the optimal full haplotype frequency *q*_*k*_ is inferred by maximizing the likelihood function:

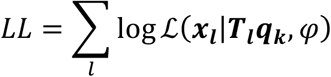

where ℒ is Dirichlet multinomial likelihood, ***x***_***l***_ is the observed variant count data for read type *l* and φ is the overdispersion parameter, assumed to be 1 × 10^−3^. Simulated annealing was used to optimise the haplotype frequencies by running two independent searches for at least 5000 states (iterations) until convergence was reached. In each state, the distribution of ***q***_***k***_was drawn from a Gaussian distribution centered at the frequency distribution of the previous state with a standard deviation of 0.05. One additional haplotype was added to the set of full haplotypes during each round of optimization.

The resulting *K* haplotypes reconstructed depend on the order in which the list of potential full haplotypes is considered. As mentioned above, paired-end reads were merged to produce longer reads (up to ∼500-600 base pairs) for mapping in the case of the seasonal A/H3N2 virus samples. Additionally, the single-stranded A/H1N1pdm09 viral reads generated from 454 sequencing can be as long as ∼500 base pairs. Consequently, there was a non-trivial number of reads where co-mutations were observed in multiple polymorphic sites. Since iSNV frequencies are generally low, haplotypes with co-mutating sites would inevitably be relegated to end of the list order if ranked by their expected joint probabilities. As such, the list of full potential haplotypes was ordered in descending order based on the score of each full haplotype set *k* (*s*_*k*_):

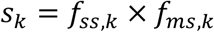

where *f*_*ss,k*_ and *f*_*ms,k*_ are both joint probabilities of the full haplotype *k* computed in different ways. *f*_*ss,k*_ is the expected joint probability frequency calculated from the observed independent frequencies of each variant for each polymorphic site found in the full haplotype *k. f*_*ms,k*_ is based on the observed frequencies of variants spanning across the sets of highest hierarchal combination of polymorphic sites (*f*_*ms,k*_).

For example, given a segment where iSNVs were found in three sites, the following reads were mapped: (A, A, C), (T, A, C), (A, T, C), (A, C, –), (–, A, C) and (–, T, C). We can immediately see that the top hierarchal combination of polymorphic sites (i.e. possible haployptes) are (A, A, C), (T, A, C) and (A, T, C) (i.e. we would compute *f*_*ms,(A,A,C)*_, and *f*_*ms,(T,A,C)*_ respectively). The observed number of reads with (–, A, C) will counted towards the computation of both *f*_*ms,(A,A,C)*_ and *f*_*ms,(T,A,C)*_ since they could be attributed to either haplotype. Similarly, reads with (–, T, C) will be absorbed towards the counts to compute *f*_*ms,(A,T,C)*_. Finally, we see that reads with (A, C, –) are not a subset of any of the top hierarchal haplotypes considered. As such, they form the 4^th^ possible top hierarchal haplotype on its own. As such, if we were to compute the ranking for haplotype (A, A, C):

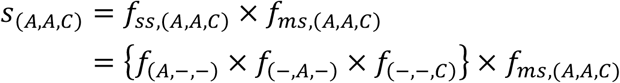

If any nucleotide variants in the observed partial haplotypes were unaccounted for in the current round of full haplotypes considered, they were assumed to be generated from a cloud of “noise” haplotypes that were present in no more than 1%. Bayesian information criterion (BIC) was computed for each set of full haplotypes considered and the most parsimonious set of haplotypes was determined by the lowest BIC value.

### Linkage disequilibrium

Using the estimated frequencies of the most parsimonious reconstructed haplotypes, conventional Lewontin’s metrics of linkage disequilibrium were computed to detect for potential epistatic pairs of nonsynonymous variants:

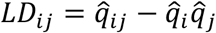

where 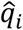 and 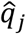 are the estimated site-independent iSNV frequencies of sites *i* and *j* respective while 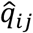 is the frequency estimate of variants encoding co-variants in both *i* and *j*. Dividing *LD* by its theoretical maximum normalises the linkage disequilibrium measure:

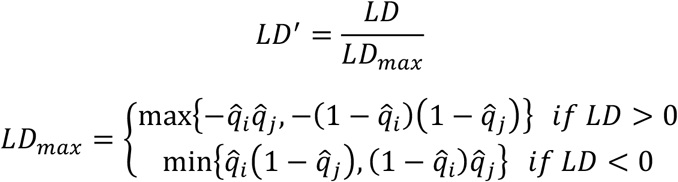

### FoldX analyses

FoldX (https://foldxsuite.crg.eu/) was used to estimate structural stability effects of likely linked nonsynonymous minority variants found in the nucleoprotein (NP) of within-host A/H3N2 virus populations. At the time of writing of this paper, there was no A/H3N2-NP structure available. Although the eventual NP structure (PDB: 3ZDP) adopted for stability analyses was originally derived from H1N1 virus (A/WSN/33)^40^, it was the most well resolved (2.69Å) crystal structure available, with 78.5% amino acid identity relative to the NP protein of A/Brisbane/10/2007. Previous work has shown that mutational effects predicted by FoldX using a NP structure belonging to A/WSN/33 (H1N1) was similar to those experimentally determined on a A/Brisbane/10/2007 nucleoprotein^31^. FoldX first removed any potential steric clashes to repair the NP structure. It then estimated differences in free energy changes as a result of the input amino acid mutation (i.e. Δ Δ *G* = Δ *G* _*mutant*_ − Δ *G*_*wild−type*_) under default settings (298K, 0.05M ionic strength and pH 7.0). Five distinct simulations were made to estimate the mean and standard deviation Δ Δ *G* values.

### Phylogenetic inference

All maximum likelihood phylogenetic trees were reconstructed with IQTREE (v. 1.6.10)^73^, using the GTR+I+G4 nucleotide substitution model.

## Data availability

All raw sequence data have been deposited at NCBI sequence read archive under BioProject Accession number PRJNA722099. All custom Python code and Jupyter notebooks to reproduce the analyses in this paper are available online: https://github.com/AMC-LAEB/Within_Host_H3vH1.

## Acknowledgements

We thank Carolien van de Sandt for helpful discussions. We gratefully acknowledge the authors, originating and submitting laboratories (Table S5) for the reference sequences retrieved from GISAID’s EpiFlu Database used in this study.

A.X.H., Z.C.F.G. and C.A.R. were supported by ERC NaviFlu (No. 818353). The South East Asia Infectious Disease Clinical Research Network (SEAICRN) was funded by National Institutes of Allergy and Infectious Diseases, National Institutes of Health (US), N01-A0-50042, HHSN272200500042C.

## Author contributions

A.X.H., Z.C.F.G., M.R.A.W., D.E., M.D.d.J., and C.A.R. designed the research; A.X.H., Z.C.F.G. and M.R.A.W. performed the data analyses; M.R.A.W., R.M.V., T.N.D, L.T.Q.M., P.Q.T., T.T.N.A., H.M.T., N.T.H., L.Q.T., L.T.H., H.T.B.N., K.C., P.P., N.V.V.C., N.M.N., D.D.T., T.T.H., H.F.L.W., P.H., A.F., H.R.V.D., D.E. and M.D.d.J. collected the clinical samples and generated the sequencing data; A.X.H., Z.C.F.G. and C.A.R. wrote the first draft of the paper. All authors contributed to the critical review and revision of the paper.

