## Supplementary materials for "Within-host evolutionary dynamics of seasonal and pandemic human influenza A viruses in young children"

*Han et al.*

*Haemagglutinin and neuraminidase minority variants in A/H3N2 virus samples*

The amino acid variants emerging in the haemagglutinin (HA) and neuraminidase (NA) proteins of A/H3N2 virus samples were discussed in a previous work[1]. Briefly, given the lack of antibody-mediated immune response in this cohort of mostly naïve children, HA amino acid variants emerging in putative antigenic sites were generally low in frequencies (median frequency = 0.04, IQR=0.03-0.06) and all only became detectable 3-4 days post symptom onset (Figure S5A). Notably, two of these intra-host mutations were also found in the global A/H3N2 virus population in high frequencies: HA-S45N and -D53N (H3 numbering without signal peptide). Both mutations are part of the canonical antigenic site C of H3 and emerged within separate individuals, with D53N eventually becoming the majority variant (97%) as late as day 13 post-symptom onset in one of them. Oseltamivir-resistant amino acid mutations E119V, R292K and N329K arose in 10 patients that were treated with the antiviral drug and mostly rose in within-host frequencies only 4-7 days after administration of oseltamivir (Figure S5B).

*Genetic diversity by $\pi$ statistic*

Given that the number of identified iSNVs can potentially be biased by variations in sequencing coverage, genetic diversity was also assessed using nucleotide diversity $\pi$ statistic[2]. This approach constitutes a more robust measure of within-host diversity as it is solely dependent of the underlying variant frequencies. While the number of polymorphic sites provides an estimate of "richness" in the viral population, it may be incompatible to compare iSNV counts between samples with different read coverage profiles[3]. In contrast, $\pi$ is a more robust metric that is unbiased by sequencing depth. For each site $l$:

$$\pi_l = \frac{N_l(N_l - 1) - \sum_j n_{j,l}(n_{j,l} - 1)}{N_l(N_l - 1)}$$

where $N_l$ and $n_{j,l}$ are the coverage and number of reads encoding allele $j$ in site $l$ respectively[3]. To compute the $\pi$ statistic for the entire genome of length $L$:

$$\pi = \sum_{l=1}^{L} \frac{\pi_l}{L}$$

Here, we observed similar trends in genetic diversity when using $\pi$ statistics compared to iSNVs counts (Figure S8). $\pi$ weakly increased with respect to time and CT values for A/H3N2 viruses (days since illness onset: Spearman $\rho = 0.388$, $p = 1.52 \times 10^{-8}$; CT: $\rho =$

0.455, $p = 3.66 \times 10^{-10}$) while remaining relatively invariant for A/H1N1pdm09 viruses (days since symptom onset: $\rho = 0.017, p = 0.92$; CT: $\rho = 0.240, p = 0.13$).

*Potential linked minority variants in within-host virus populations*

For both within-host seasonal A/H3N2 and pandemic A/H1N1 virus populations, there were few instances of potentially linked nonsynonymous variants and if such co-variants were to exist, they were mainly found in the internal gene segments (Table S2). There was only one pair of HA amino acid mutations (E261G/L455F) that was encoded by a minority haplotype of A/H1N1pdm09 viruses infecting one individual but the normalized Lewontin's linkage disequilibrium measure ($LD'$) was less than 0.5, suggesting a low likelihood that the mutation pair was linked non-randomly. These potentially linked variants tend to emerge late in the infection (6-7 days post illness onset) for both viral subtypes, in inferred haplotypes appearing at low frequencies within-host (median frequency = 0.08, IQR = 0.03-0.12) that were not shared between multiple individuals.

*Binomial sampling models for transmission bottlecneck estimation*

Index cases were previously identified for six of the seven households where the pandemic A/H1N1pdm09 viral samples were collected[4]. Assuming that the non-index cases within the same household were secondarily infected by the index case, six transmission pairs were identified where samples with adequate breadth of coverage (>70% of genome covered with >50x coverage) were collected from the index patient on an earlier date relative to the secondary case.

Virus transmission bottleneck sizes were then estimated using two binomial sampling models that were elaborated in detail by [5] and [6]. First, the presence/absence model computes transmission probability as the probability that a transmitted donor iSNV was found in at least one genome in the bottleneck population:

$$P_{d,i}(A|N_b) = p_{d,A}^{N_b}$$

where $A$ refers to the transmitted iSNV in polymorphic site $i$, $N_b$ is the bottleneck size and $p_{d,A}$ is the frequency of allele $A$ in the sampled virus population within donor $d$.

The presence/absence model does not incorporate recipient frequencies of transmitted iSNVs in its probability calculations. It assumes that all transmitted iSNVs are detected in the recipient, and thus any donor iSNVs that are not present in the recipient are considered to have not been transmitted. It also does not account for any changes to $p_d$ between the time of sampling and day of transmission. The maximum likelihood estimate of $N_b$ would thus yield the largest log likelihood value given by:

$$LL(N_b) = \sum_d \sum_i \ln P_{d,i}$$

To incorporate information on recipient frequencies which can change between transmission and sampling, transmission bottleneck sizes were re-estimated using a second beta-binomial model formulated by Sobel Leonard et al. (2017). For each allele $A$ observed in polymorphic site $i$ that was transmitted from donor $d$ to recipient $r$, the log-likelihood of $N_b$ is given as:

$$LL(N_b)_{d,r}^{transmitted} = \sum_{A_i} \ln \left\{ \sum_{k=1}^{N_b} p_{beta}\left(p_{r,A_i} \middle| k, N_b - k\right) p_{bin}\left(k \middle| N_b, p_{d,A_i}\right) \right\}$$

where $p_{beta}\left(p_{r,A_i} \middle| k, N_b - k\right)$ is the conditional probability density, as modelled by the beta distribution, that the transmitted iSNV, $A_i$ is found in the recipient at frequency $p_{r,A_i}$ given that the variant is found present in $k$ genomes out of the total transmission bottleneck of $N_b$ genomes. $p_{bin}\left(k \middle| N_b, p_{d,A_i}\right)$ is the binomial probability of drawing $k$ genomes with allele $A_i$ in a sample of $N_b$ genomes and variant frequency of $p_{d,A_i}$ within the donor.

As some of the iSNVs in the donor may not be transmitted or were present below the 2% minimum variant frequency cut-off, the likelihood of these events was computed by:

$$LL(N_b)_{d,r}^{lost} = \sum_{A_i} \ln \left\{ \sum_{k=1}^{N_b} p_{beta,cdf}\left(p_{r,A_i} < 0.02 \middle| k, N_b - k\right) p_{bin}\left(k \middle| N_b, p_{d,A_i}\right) \right\}$$

where $p_{beta,cdf}$ is the cumulative distribution function of the beta distribution.

The maximum likelihood estimate of $N_b$ as described by the beta-binomial was then computed by searching for the value of $N_b$ that would give the largest value of:

$$LL(N_b) = \sum_{d,r} LL(N_b)_{d,r}^{transmitted} + LL(N_b)_{d,r}^{lost}$$

Log-likelihood values were computed for a range $N_b$ between 1 to 1000 genomes for both models.

*Transmission bottleneck size estimation of pandemic A/H1N1pdm09 viral infections*

Index cases were epidemiologically inferred for four of the seven households where pandemic A/H1N1pdm09 virus samples were collected[4]. Assuming all other patients in the same household were secondary cases infected by the index patient, five transmission pairs were identified (Figure S13A). Adding further support that the inferred transmission pairs were likely linked by true transmission events, the per-nucleotide site L1-norm genetic distance distribution of the four transmission pairs identified (median L1-norm distance = $6.63 \times 10^{-3}$ divergence per site, IQR = $6.04 \times 10^{-3} - 7.57 \times 10^{-3}$ divergence per site) is significantly lower than the genetic distances of sample pairs collected from different individuals in the same community where these households were located, on the same day since the patients' respective symptom onset date (median L1-norm distance = 0.024 divergence per site, IQR = 0.020-0.029 divergence per-site; Mann-Whitney U $p$-value = 0.02; Figure S13B).

Transmission bottleneck size was then estimated by aggregating over all transmission pairs and applying two previously developed sampling based models[5,6]. We estimated the transmission bottleneck of pandemic A/H1N1pdm09 viruses to be 1-2 genomes (maximum likelihood (ML) presence-absence model estimate = 1 genome; ML beta-binomial model estimate = 2 genomes). The tight bottleneck was further reflected in the iSNV frequency plot where most of the donor variants were either present as the single majority allele or were not transmitted/undetected in the recipient, with few shared iSNVs between the two virus populations (Figure S13A). Furthermore, we observed that the virus populations between patients (median L1-norm distance = 0.031 divergence per site, IQR = 0.024-0.046 divergence per site) were significantly more different than longitudinal samples collected from the same individual (median L1-norm distance = $2.35 \times 10^{-3}$ divergence per site, IQR = $1.54 \times 10^{-3} - 3.18 \times 10^{-3}$ divergence per site), suggesting that the haplotypes found within an individual remains relatively invariant throughout the course of infection (Figure S13B). Combined with the fact that the per-site L1-norm genetic distance between samples attributed to the identified transmission pairs (median L1-norm distance = $6.49 \times 10^{-3}$ divergence per site, IQR = $6.34 \times 10^{-3} - 6.63 \times 10^{-3}$ divergence per site) was greater than those computed for longitudinal samples pairs of each household individual (median L1-norm distance = $2.77 \times 10^{-3}$ divergence per site, IQR = $2.18 \times 10^{-3} - 3.32 \times 10^{-3}$ divergence per site; Mann-Whitney U $p$-value = 0.01; Figure S13B), it is likely only a limited number of haplotypes were shared between individuals in a transmission pair. Based on the most parsimonious reconstructed haplotypes for the five transmission pairs encoding shared iSNVs, we estimated the median number of haplotypes transmitted from donor to recipient to be between 1 and 2 haplotypes.

*Mutation-selection balance*

Considering a single-locus mutant with deleterious fitness effect $s$ (i.e. $s < 0$), the frequency of the mutant allele ($f$) can be modelled by the following stochastic differential equation, otherwise known as the Langevin equation[7]:

$$\frac{\partial f}{\partial t} = \underbrace{sf(1-f)}_{\text{selection}} + \underbrace{\sqrt{\frac{f-(1-f)}{N}}\eta(t)}_{\text{genetic drift}} + \underbrace{\mu(1-f)}_{\text{mutation}}$$

where $N$ is the population size, $\mu$ is the mutation rate and $\eta(t)$ is the stochastic noise term due to genetic drift. For any Langevin equation, we can find the time-dependent probability distribution of $f$ (i.e. $\frac{\partial p(f,t)}{\partial t}$) by its corresponding Fokker-Planck equation. At stationarity (i.e. $\frac{\partial p(f,t)}{\partial t} = 0$), its solution is known to be:

$$p(f) \propto \frac{e^{-2N\Lambda(f)}}{f(1-f)}$$

where $-\frac{\partial \Lambda(f)}{\partial t} = s + \frac{\mu}{f}$

Integrating $-\frac{\partial \Lambda(f)}{\partial t}$, we will get:

$$-\Lambda(f) = sf + \mu \ln f$$

which can then be substituted to obtain:

$$p(f) \propto e^{2Nsf} \cdot f^{2N\mu - 1}$$

In other words, $p(f)$ strongly depends on $N\mu$. If $N\mu \gg 1$, we can see that $p(f)$ will strongly peak at some characteristic value of $f$ that minimizes $\Lambda(f)$. If $N$ is large, we can assume drift effects is negligible and $f$ is largely deterministic due to selection:

$$\frac{\partial f}{\partial t} = sf(1-f) + \mu(1-f) = \left\{-\frac{\partial \Lambda(f)}{\partial t}\right\}[f(1-f)]$$

$$\Rightarrow \frac{\partial \Lambda}{\partial t} = \frac{\partial f}{\partial t}\left(\frac{\partial \Lambda}{\partial f}\right) = \frac{\partial f}{\partial t}\left\{-\frac{\partial f}{\partial t}\left(\frac{1}{f(1-f)}\right)\right\} \leq 0$$

In other words, selection dynamics minimises the $\Lambda(f)$ term and as such, result in mutation-selection balance:

$$-\frac{\partial \Lambda(f)}{\partial t} = s + \frac{\mu}{f} = 0$$

$$\Rightarrow f = -\frac{\mu}{s}$$

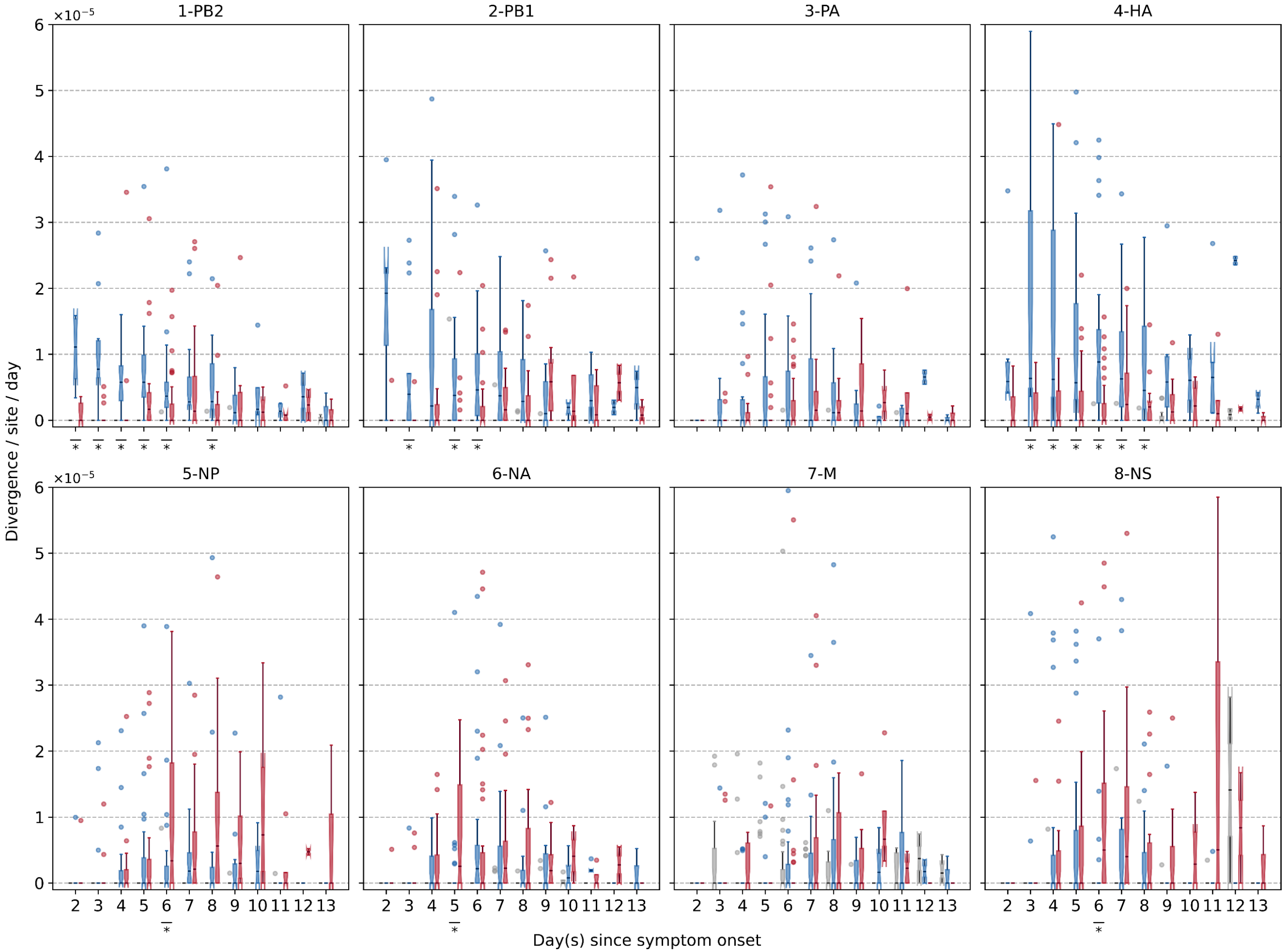


**Figure S1**: Box plots (median, interquartile range (IQR), and whiskers extending within median $\pm 1.5 \times$ IQR) summarising the empirical within-host evolutionary rates ($r_{g,t}$) of different H3N2 viral gene segments. All rates are stratified by substitution type (synonymous – blue; nonsynonymous – red; stop codon – grey). Wilcoxon signed-rank tests were performed to assess if the paired synonymous and nonsynonymous evolutionary rates are significantly distinct per timepoint (annotated with "$*$" if $p < 0.05$).

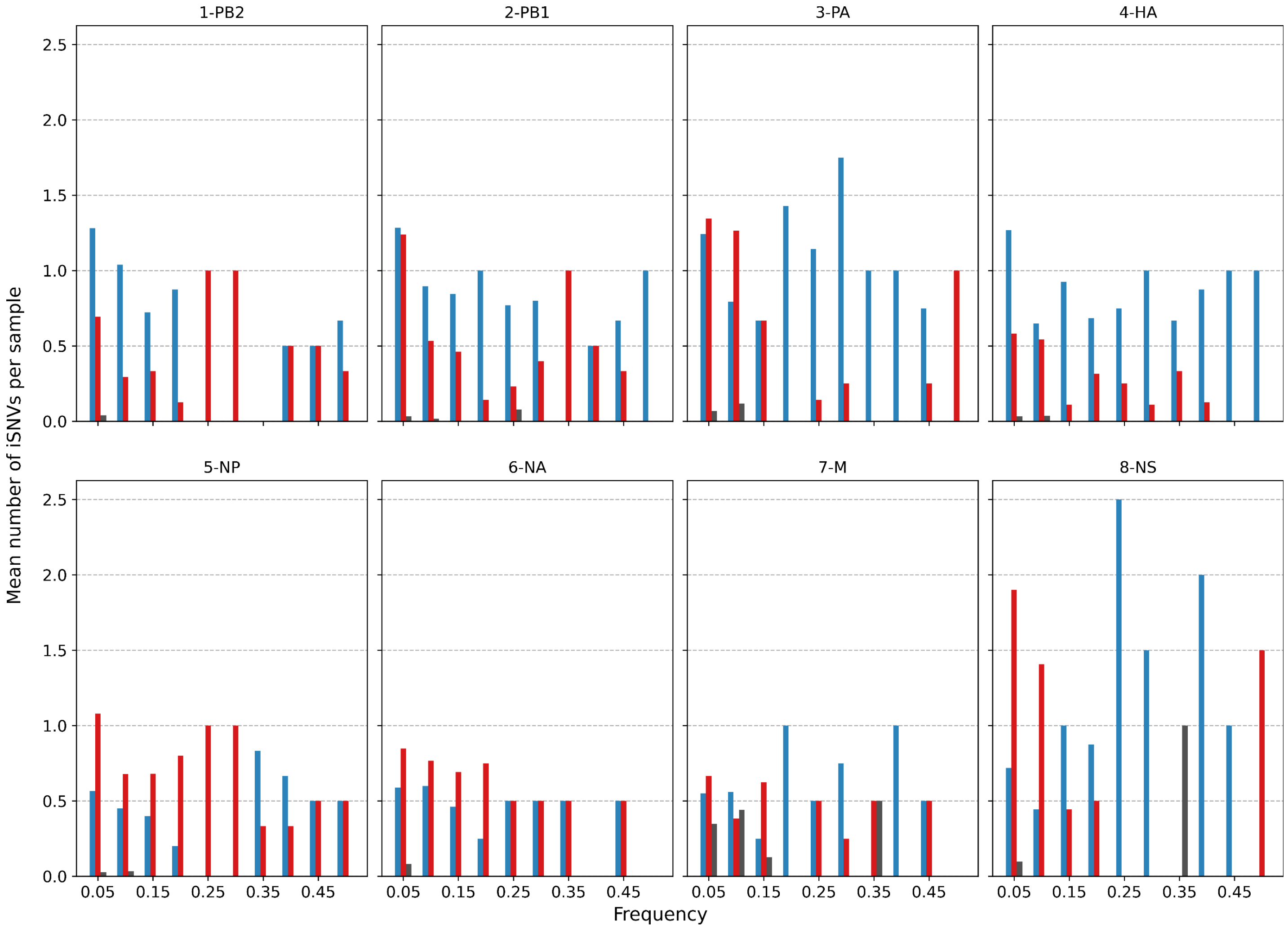


**Figure S2**: Histogram of the mean number of minority iSNVs identified per sample across all H3N2 viral gene segments across all samples sorted by frequency bins of 5% and substitution type (synonymous – blue; nonsynonymous – red; grey – stop codon).

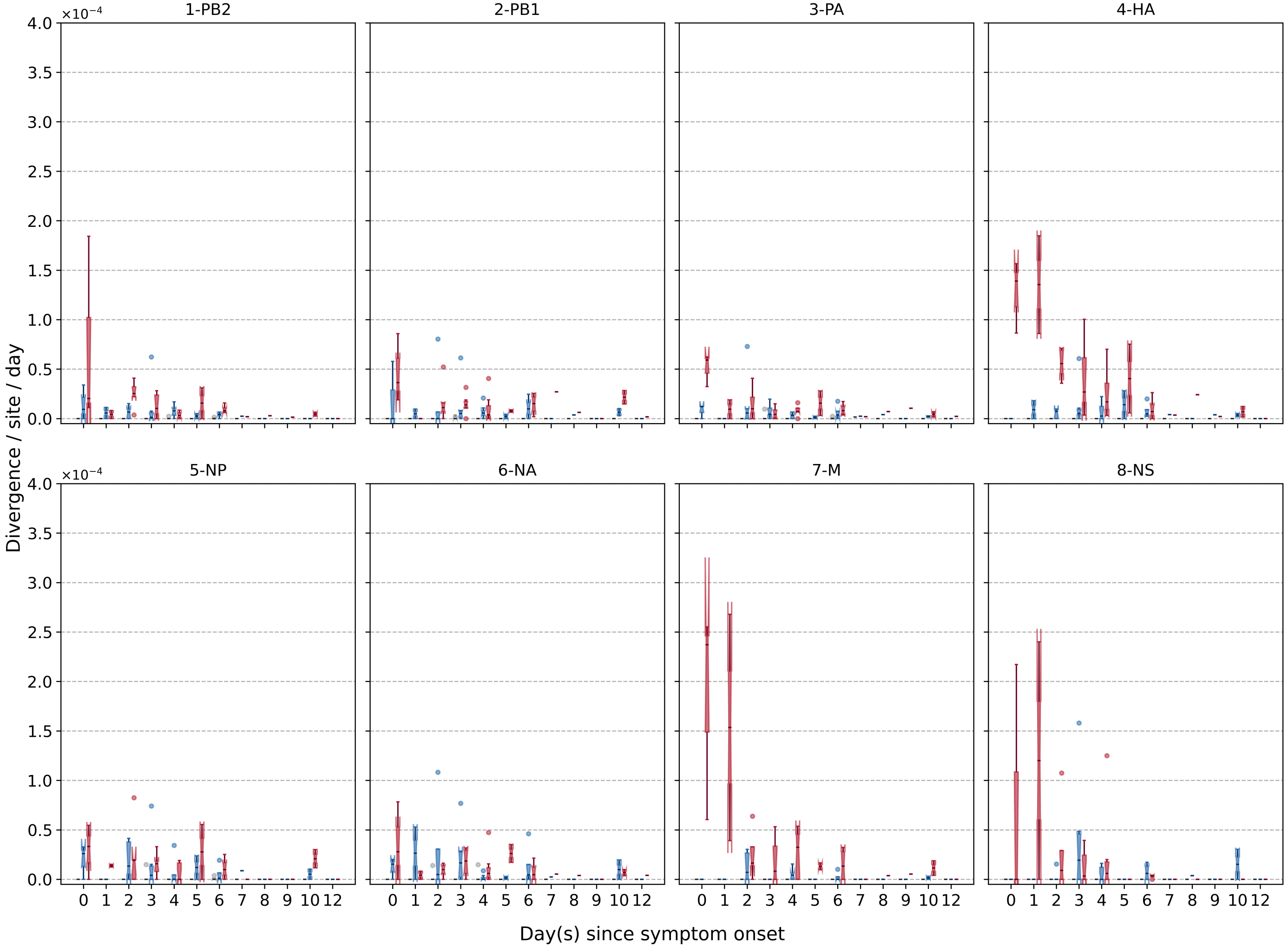


**Figure S3**: Box plots (median, interquartile range (IQR), and whiskers extending within median $\pm 1.5 \times$ IQR) summarising the empirical within-host evolutionary rates ($r_{g,t}$) of different H1N1pdm09 viral gene segments. All rates are stratified by substitution type (synonymous – blue; nonsynonymous – red; stop codon – grey). Wilcoxon signed-rank test was *not* performed here due to low number of samples collected (i.e. median number of samples per day post illness onset = 2).

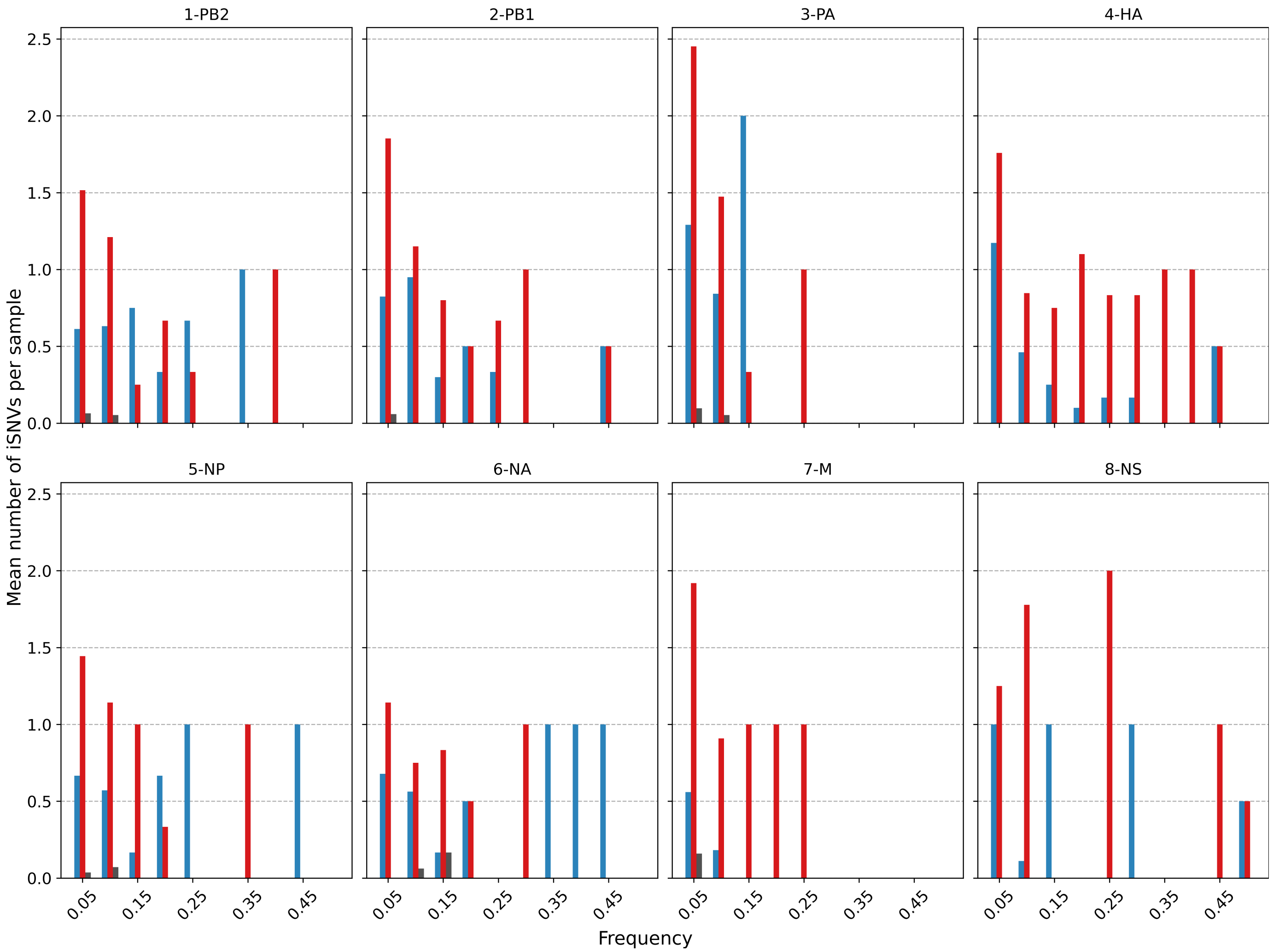


**Figure S4**: Histogram of the mean number of minority iSNVs identified across all H1N1pdm09 viral gene segments across all samples sorted by frequency bins of 5% and substitution type (synonymous – blue; nonsynonymous – red; stop-codon – grey).

A

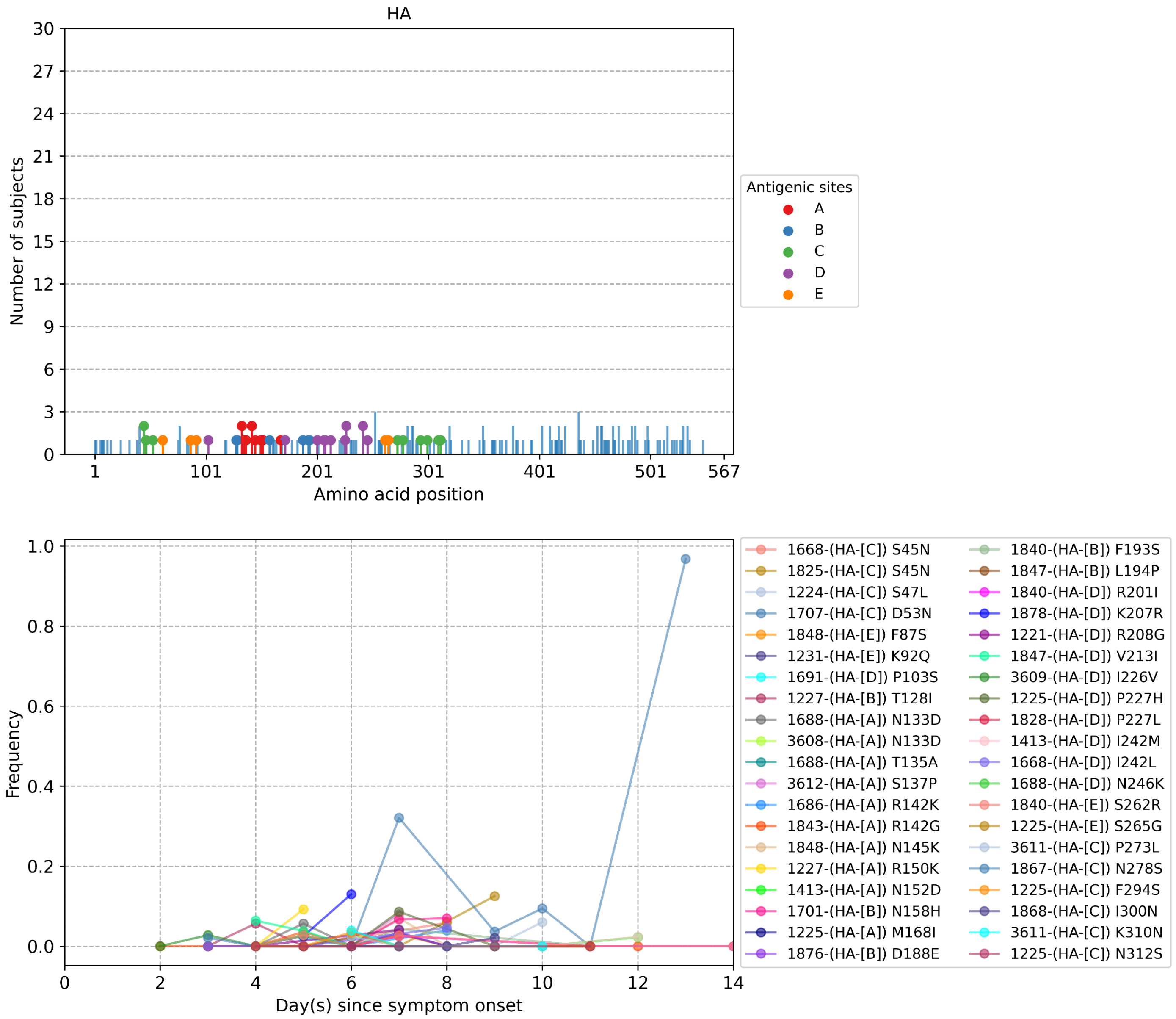

HA
Number of subjects
Amino acid position
Antigenic sites
A
B
C
D
E
Frequency
Day(s) since symptom onset
1668-(HA-[C]) S45N
1825-(HA-[C]) S45N
1224-(HA-[C]) S47L
1707-(HA-[C]) D53N
1848-(HA-[E]) F87S
1231-(HA-[E]) K92Q
1691-(HA-[D]) P103S
1227-(HA-[B]) T128I
1688-(HA-[A]) N133D
3608-(HA-[A]) N133D
1688-(HA-[A]) T135A
3612-(HA-[A]) S137P
1686-(HA-[A]) R142K
1843-(HA-[A]) R142G
1848-(HA-[A]) N145K
1227-(HA-[A]) R150K
1413-(HA-[A]) N152D
1701-(HA-[B]) N158H
1225-(HA-[A]) M168I
1876-(HA-[B]) D188E
1840-(HA-[B]) F193S
1847-(HA-[B]) L194P
1840-(HA-[D]) R201I
1878-(HA-[D]) K207R
1221-(HA-[D]) R208G
1847-(HA-[D]) V213I
3609-(HA-[D]) I226V
1225-(HA-[D]) P227H
1828-(HA-[D]) P227L
1413-(HA-[D]) I242M
1668-(HA-[D]) I242L
1688-(HA-[D]) N246K
1840-(HA-[E]) S262R
1225-(HA-[E]) S265G
3611-(HA-[C]) P273L
1867-(HA-[C]) N278S
1225-(HA-[C]) F294S
1868-(HA-[C]) I300N
3611-(HA-[C]) K310N
1225-(HA-[C]) N312S

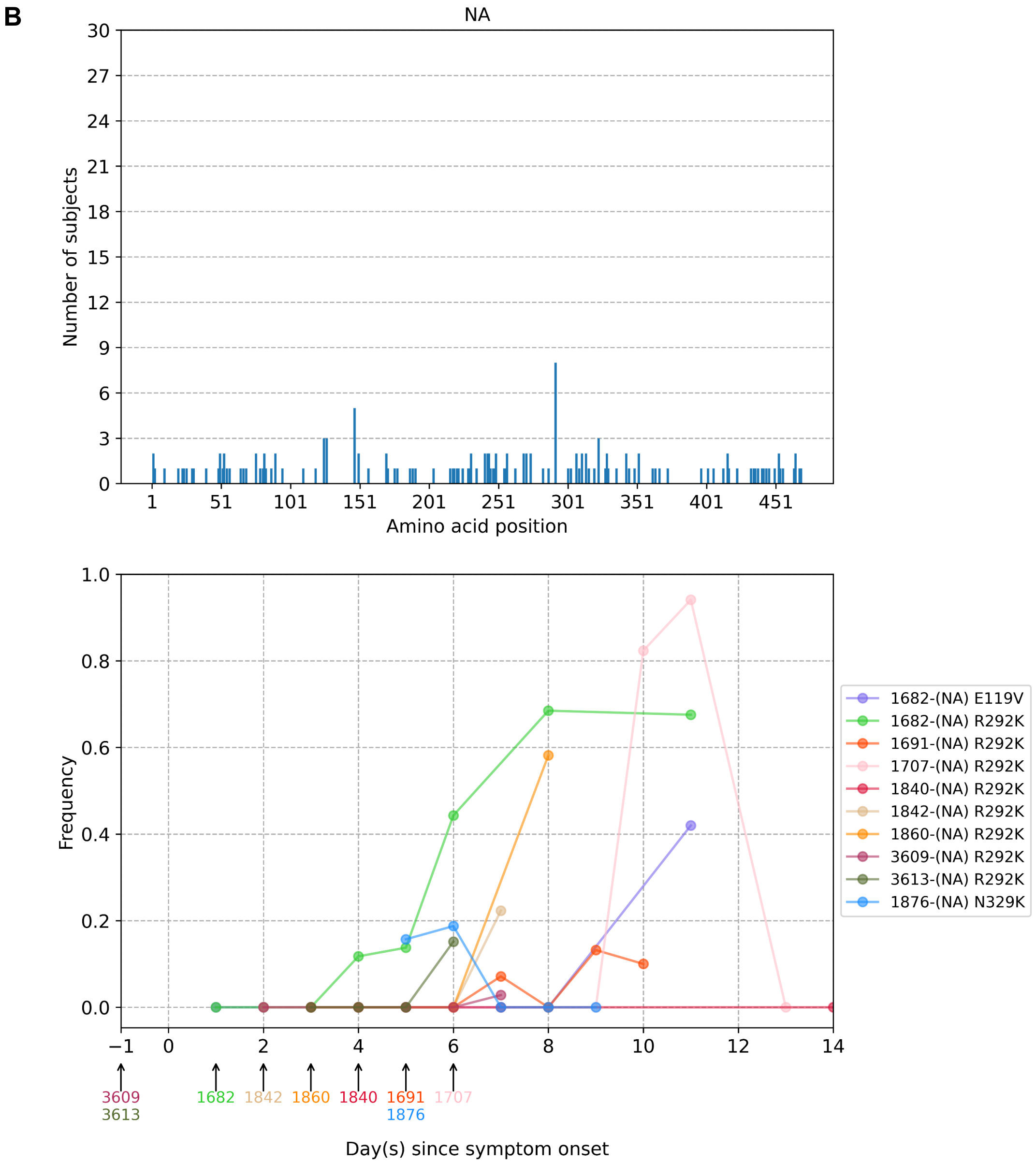
B
NA
Number of subjects
Amino acid position
Frequency
Day(s) since symptom onset
1682-(NA) E119V
1682-(NA) R292K
1691-(NA) R292K
1707-(NA) R292K
1840-(NA) R292K
1842-(NA) R292K
1860-(NA) R292K
3609-(NA) R292K
3613-(NA) R292K
1876-(NA) N329K
3609
3613
1682
1842
1860
1840
1691
1876
1707

C

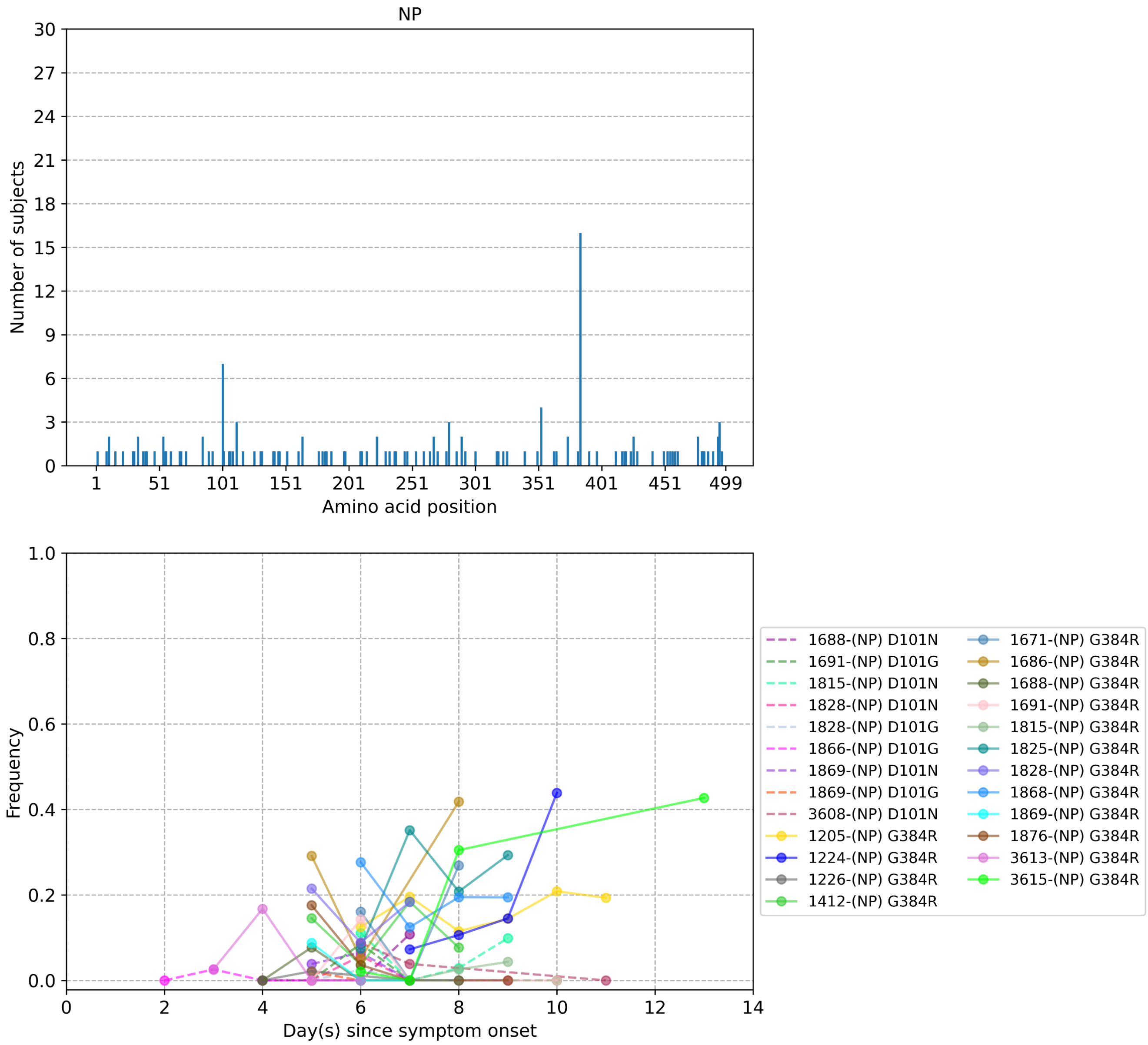
NP
Number of subjects
Amino acid position
30
27
24
21
18
15
12
9
6
3
0
1
51
101
151
201
251
301
351
401
451
499
Frequency
Day(s) since symptom onset
1.0
0.8
0.6
0.4
0.2
0.0
0
2
4
6
8
10
12
14
1688-(NP) D101N
1691-(NP) D101G
1815-(NP) D101N
1828-(NP) D101N
1828-(NP) D101G
1866-(NP) D101G
1869-(NP) D101N
1869-(NP) D101G
3608-(NP) D101N
1205-(NP) G384R
1224-(NP) G384R
1226-(NP) G384R
1412-(NP) G384R
1671-(NP) G384R
1686-(NP) G384R
1688-(NP) G384R
1691-(NP) G384R
1815-(NP) G384R
1825-(NP) G384R
1828-(NP) G384R
1868-(NP) G384R
1869-(NP) G384R
1876-(NP) G384R
3613-(NP) G384R
3615-(NP) G384R

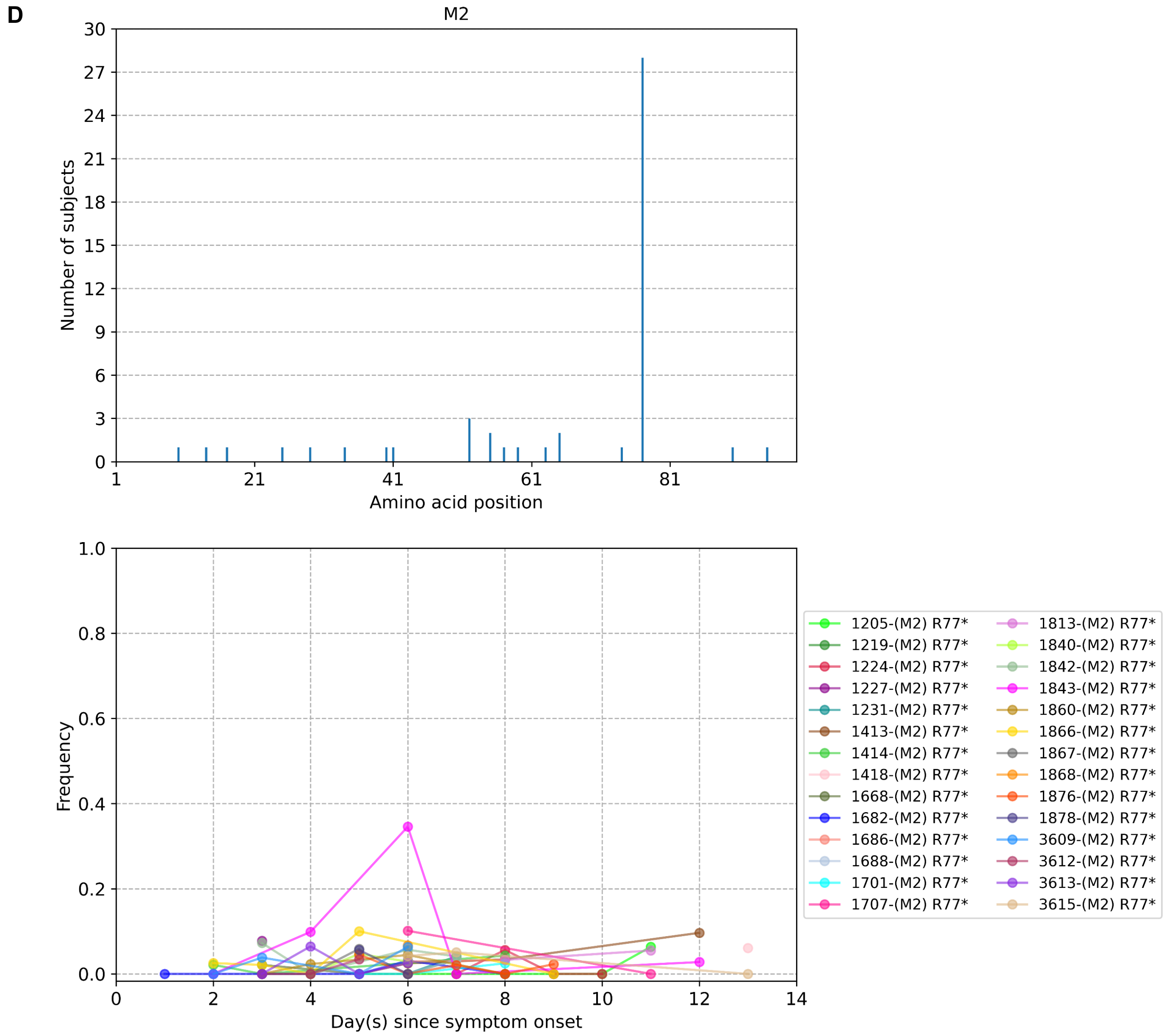
D
M2
Number of subjects
Amino acid position
Frequency
Day(s) since symptom onset
1205-(M2) R77*
1219-(M2) R77*
1224-(M2) R77*
1227-(M2) R77*
1231-(M2) R77*
1413-(M2) R77*
1414-(M2) R77*
1418-(M2) R77*
1668-(M2) R77*
1682-(M2) R77*
1686-(M2) R77*
1688-(M2) R77*
1701-(M2) R77*
1707-(M2) R77*
1813-(M2) R77*
1840-(M2) R77*
1842-(M2) R77*
1843-(M2) R77*
1860-(M2) R77*
1866-(M2) R77*
1867-(M2) R77*
1868-(M2) R77*
1876-(M2) R77*
1878-(M2) R77*
3609-(M2) R77*
3612-(M2) R77*
3613-(M2) R77*
3615-(M2) R77*

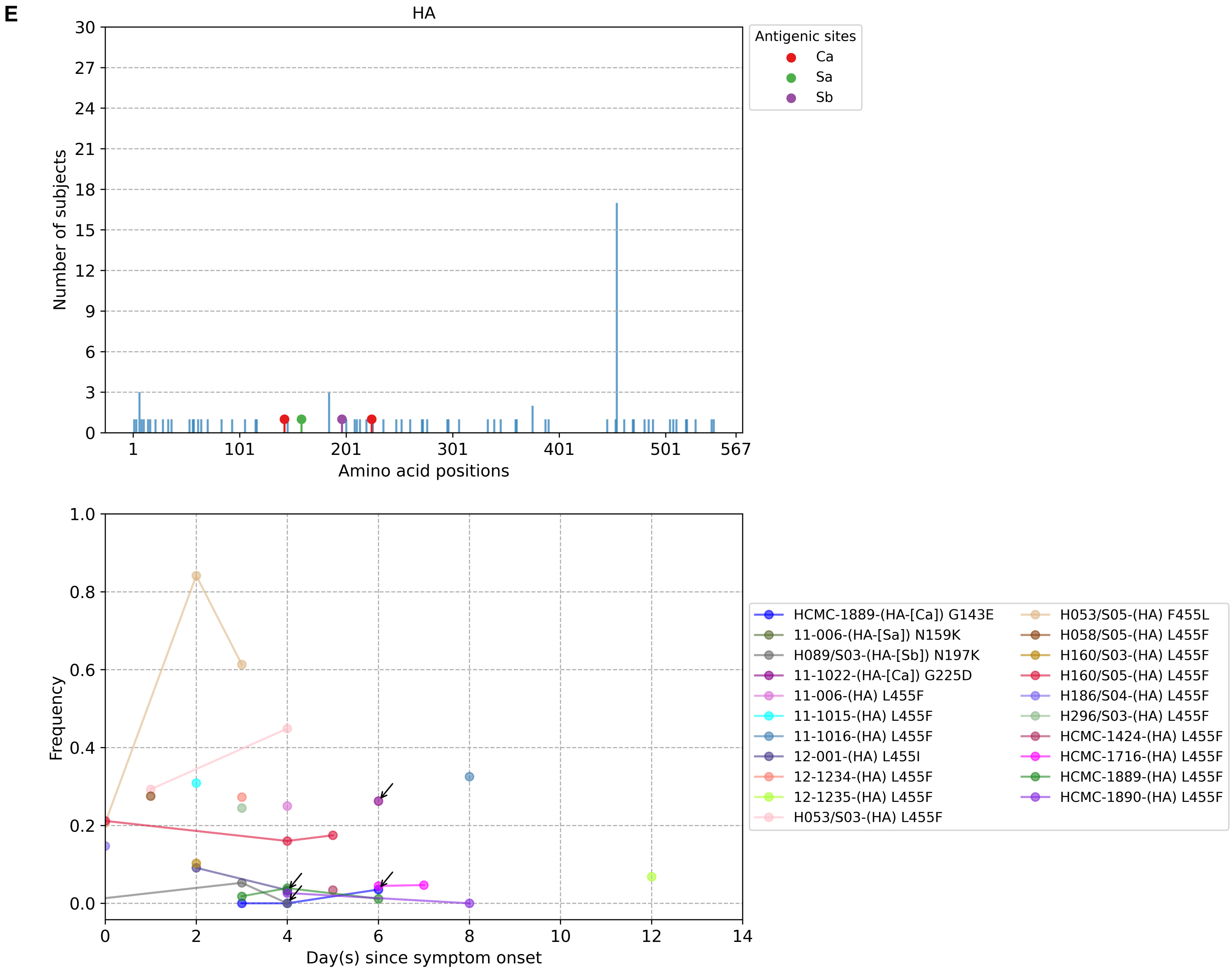

E
HA
Antigenic sites
Ca
Sa
Sb
Number of subjects
Amino acid positions
Frequency
Day(s) since symptom onset
HCMC-1889-(HA-[Ca]) G143E
11-006-(HA-[Sa]) N159K
H089/S03-(HA-[Sb]) N197K
11-1022-(HA-[Ca]) G225D
11-006-(HA) L455F
11-1015-(HA) L455F
11-1016-(HA) L455F
12-001-(HA) L455I
12-1234-(HA) L455F
12-1235-(HA) L455F
H053/S03-(HA) L455F
H053/S05-(HA) F455L
H058/S05-(HA) L455F
H160/S03-(HA) L455F
H160/S05-(HA) L455F
H186/S04-(HA) L455F
H296/S03-(HA) L455F
HCMC-1424-(HA) L455F
HCMC-1716-(HA) L455F
HCMC-1889-(HA) L455F
HCMC-1890-(HA) L455F

**F**

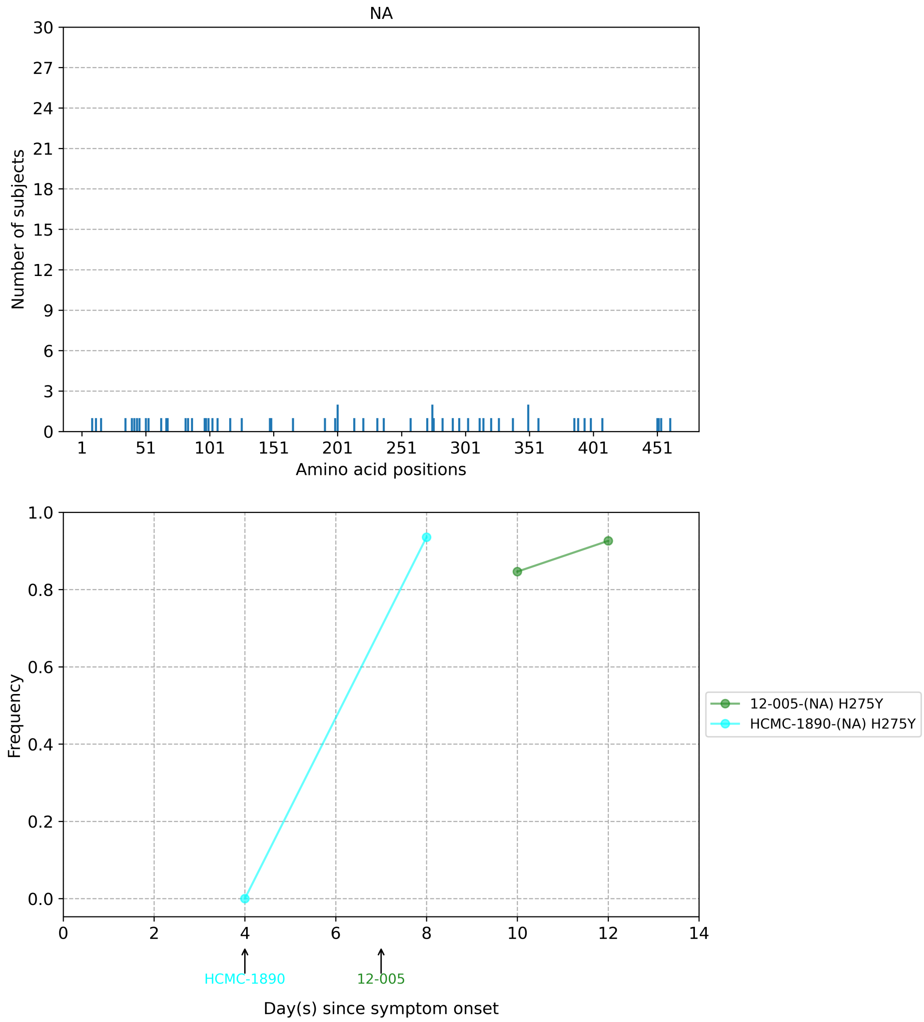
NA
Number of subjects
30
27
24
21
18
15
12
9
6
3
0
1
51
101
151
201
251
301
351
401
451
Amino acid positions
Frequency
1.0
0.8
0.6
0.4
0.2
0.0
0
2
4
6
8
10
12
14
HCMC-1890
12-005
Day(s) since symptom onset
12-005-(NA) H275Y
HCMC-1890-(NA) H275Y

**G**

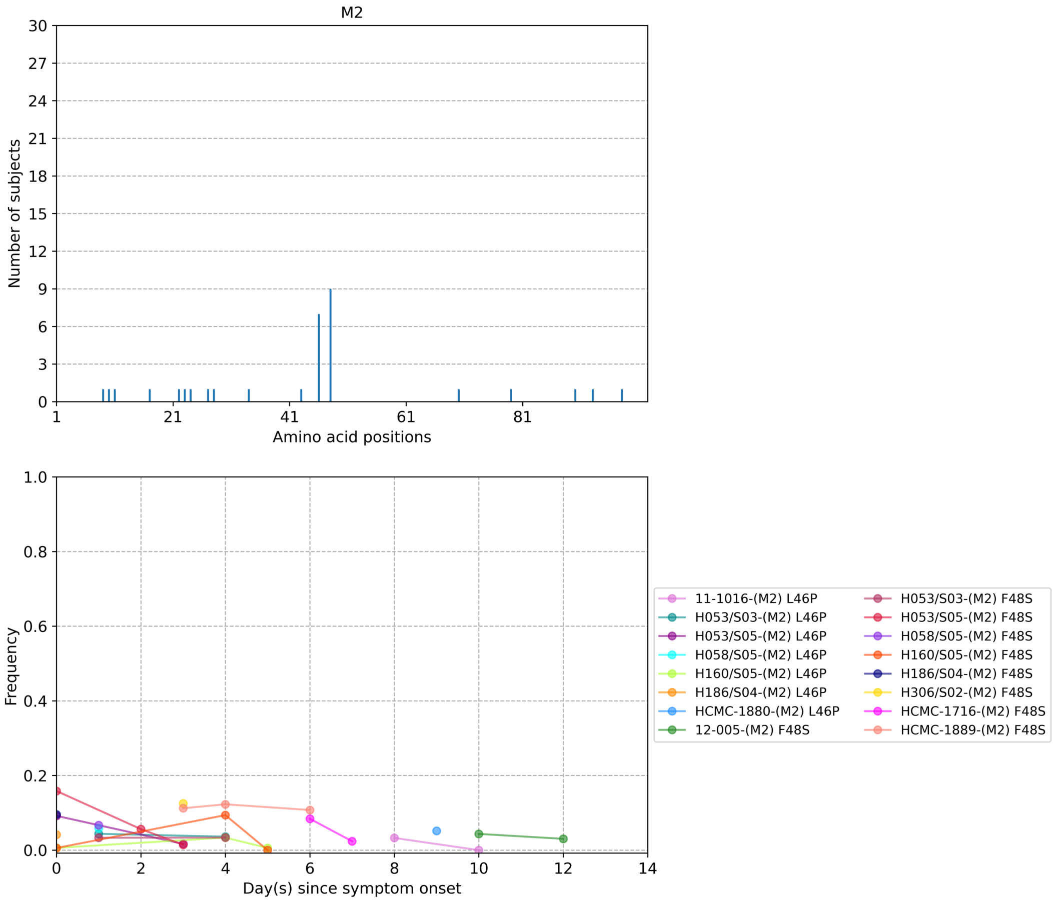


**Figure S5**: Plots of intra-host amino acid variants. Top panel shows the number of subjects where nonsynonymous variants were found in the respective protein site. Different canonical antigenic sites of the haemagglutinin (HA) protein are colored (HA numbering based on H3 numbering without signal peptide). Bottom panel plots selected as well as parallel amino acid mutations found in multiple patients against days since illness onset. Filled circles represent days on which samples were collected and sequenced. For the HA protein, the variant frequencies of all putative antigenic sites are also plotted. For the neuraminidase (NA) plot, the first day of oseltamivir treatment for individuals with resistance mutations is annotated below the *x*-axis. (**A**) HA, (**B**) NA, (**C**) nucleoprotein (NP) and (**D**) M2 protein of H3N2 viruses. (**E**) HA protein and (**F**) M2 ion channel of H1N1pdm09 viruses. The frequencies of the five putative HA antigenic variants of A/H1N1pdm09 viruses are marked by arrows for better clarity.

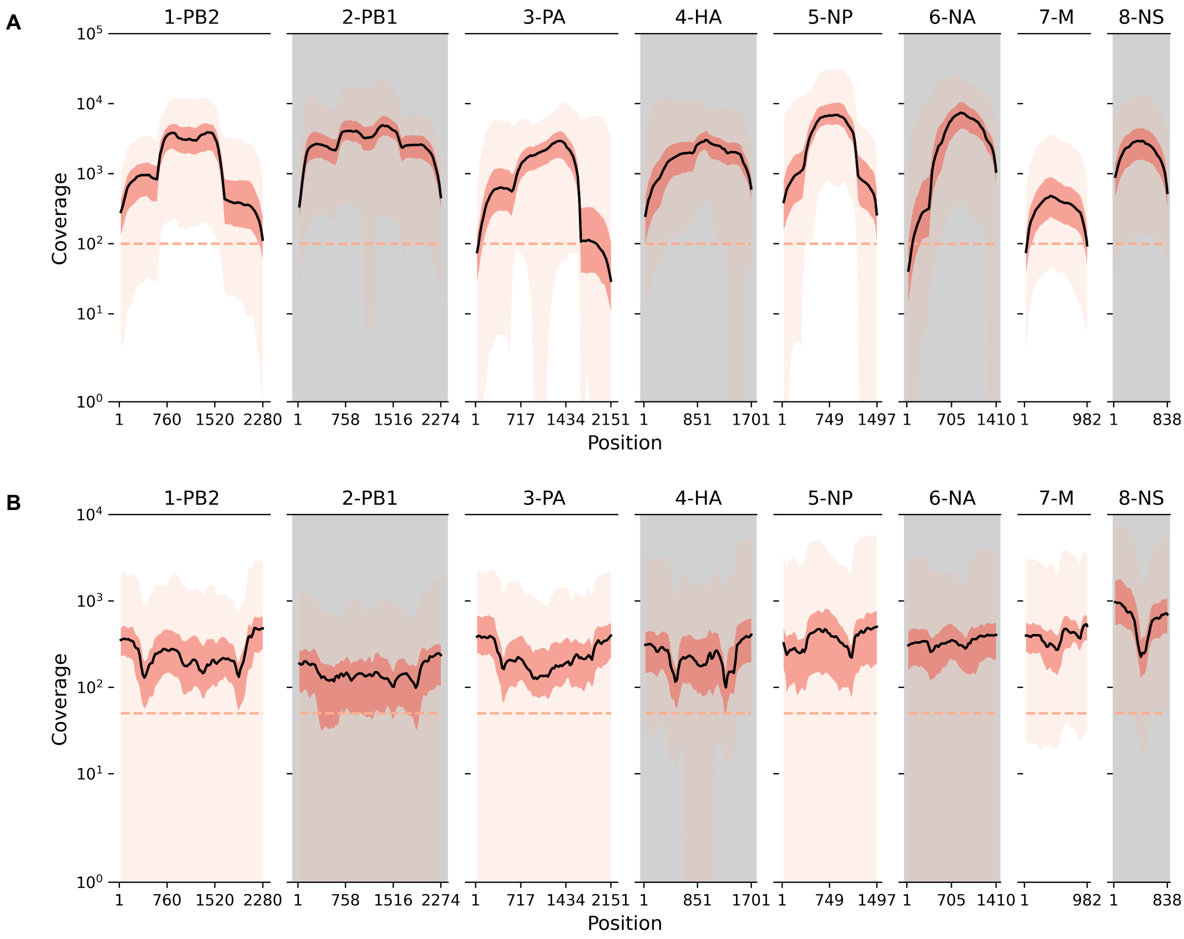


**Figure S6**: Sequence coverage across all influenza gene segments and samples. Black line plots the mean coverage for a sliding window of 50 base pairs (stepsize = 25 base pairs). The interquartile range is shaded in dark pink while the full range is denoted in light pink. (**A**) H3N2. (**B**) H1N1pdm09

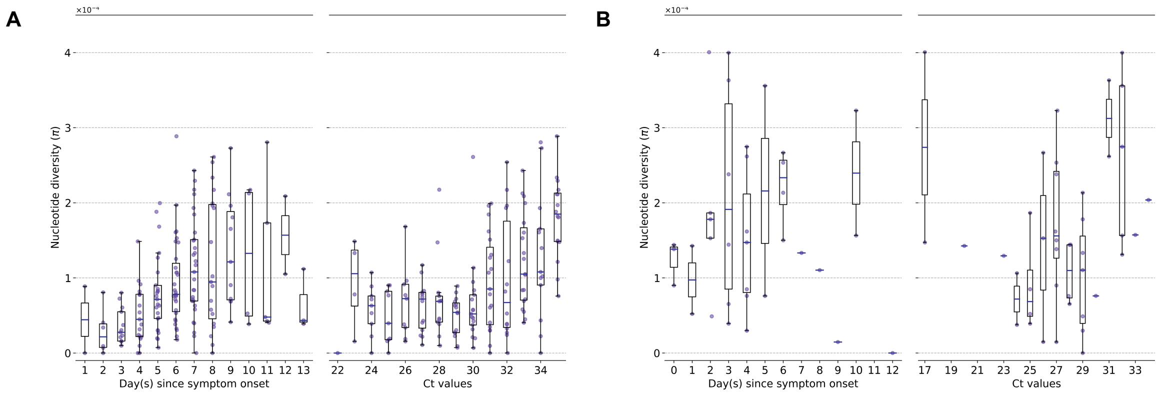


**Figure S7**: Genetic diversity of within-host influenza A virus populations as estimated by nucleotide diversity $\pi$ statistic. Box plots summarizing the $\pi$ statistic (iSNVs; median, interquartile range (IQR), and whiskers extending within median $\pm 1.5 \times$ IQR) computed for samples with adequate breadth of coverage across the whole influenza genome. (**A**) seasonal A/H3N2 and (**B**) pandemic A/H1N1pdm09 viruses. All box plots are either stratified by day(s) since symptom onset or qPCR cycle threshold (Ct) values.

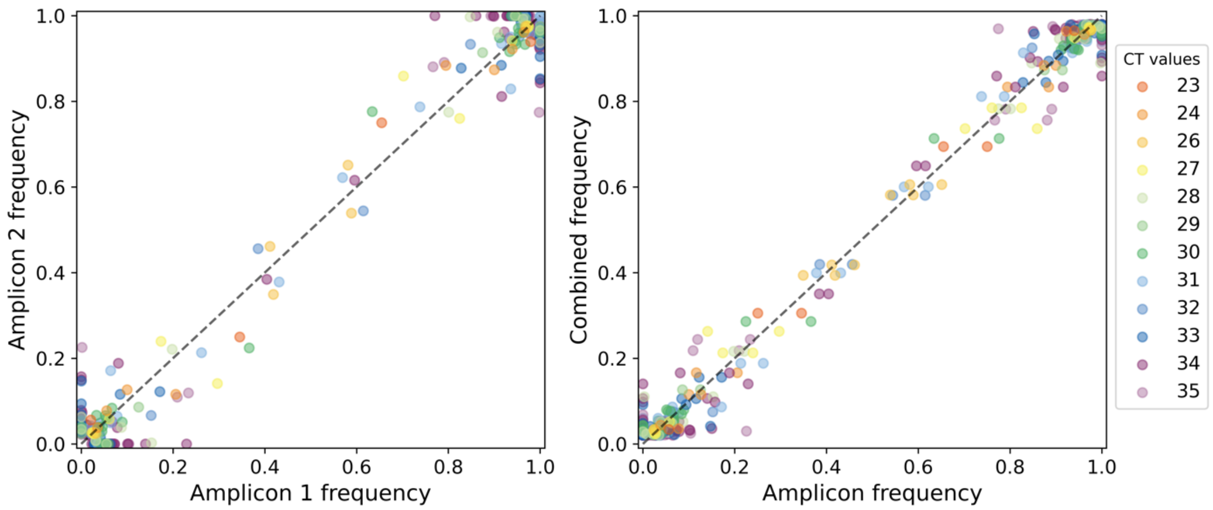


**Figure S8**: Frequencies of nucleotide variants found in A/H3N2 viral reads sequenced from overlapping amplicons. Each circle represents a nucleotide variant site (with frequency estimated between 0.02 and 0.98) found in reads attributed to at least two different amplicons (at least 100x coverage for each amplicon), and is colored by the cycle threshold (CT) value of the sample from which the variant was found. Scatter plot on the left panel compares the variant frequencies between any two amplicons while the plot on the right panel compares the variant frequencies of each amplicon to that when combining across all overlapping amplicons (i.e. the frequencies used for main analyses). The dashed line is the one-to-one expected value.

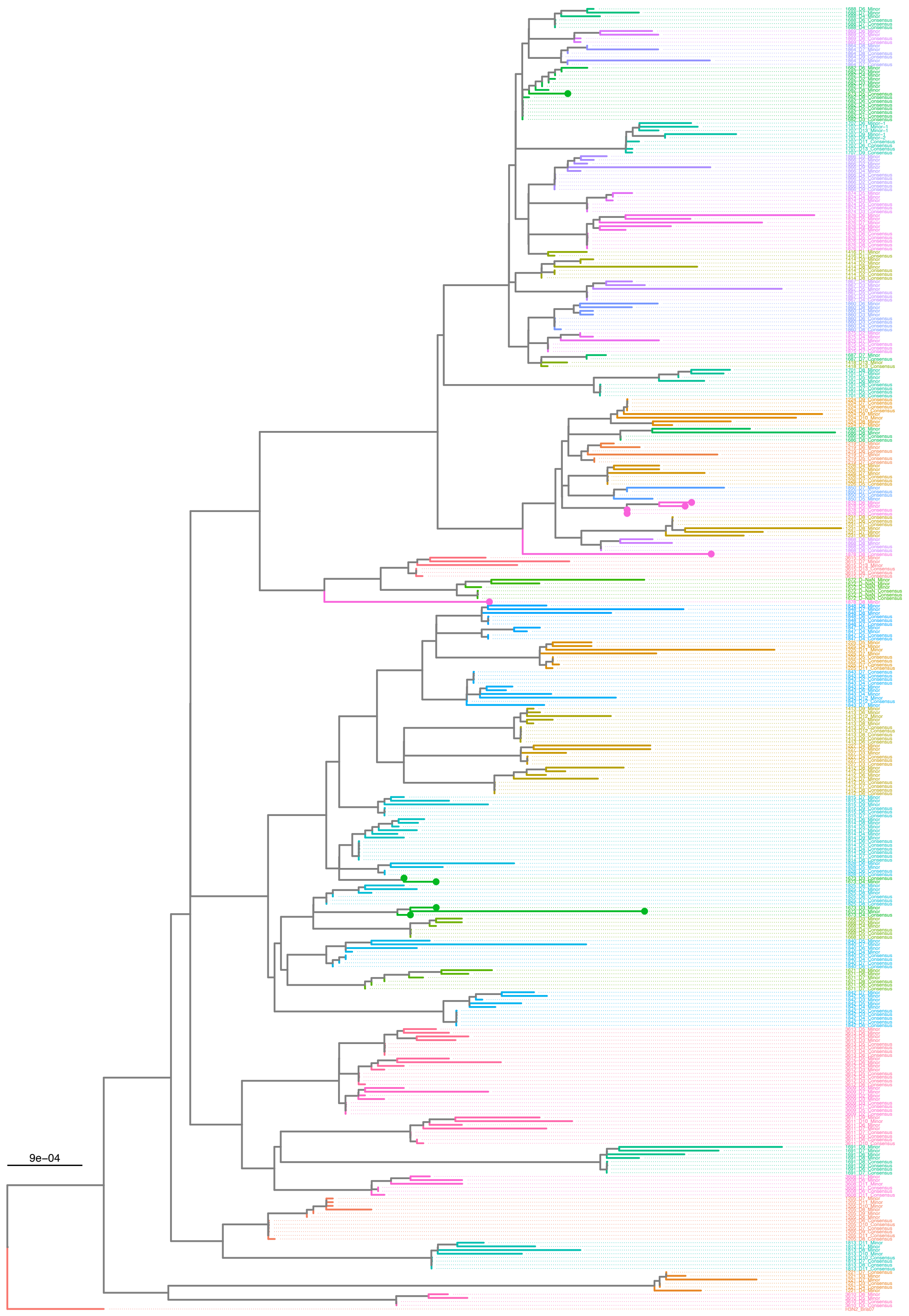


**Figure S9**: Maximum-likelihood phylogeny of putative majority (consensus) and minority whole genome sequences (by concatenating all eight gene segments) of A/H3N2 virus samples. Tip names are given in the format: “Patient ID_Days since symptom onset_putative consensus or minority sequence”. The tree is rooted to the A/Brisbane/10/2007 virus (H3N2_Bris07; EPI_ISL_103644). Subject 1673 (green tips) and the D8 sample of subject 1878 (pink tips) might have arose from mixed infections or were contaminated by other strains.

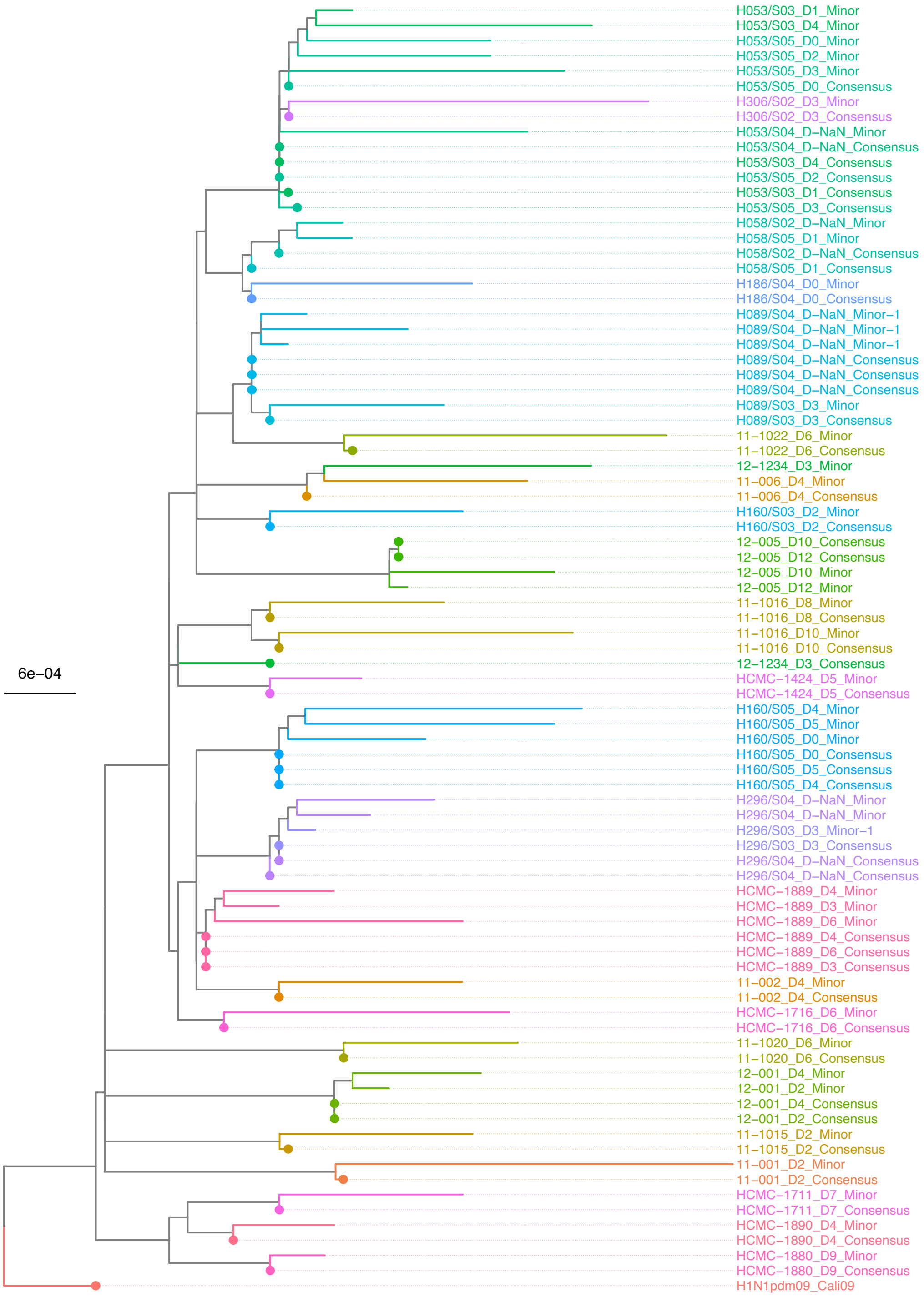


**Figure S10**: Maximum-likelihood phylogeny of putative majority (consensus) and minority whole genome sequences (by concatenating all eight gene segments) of H1N1pdm09 virus samples. Tip names are given in the format: "Patient ID_Days since symptom onset_putative consensus or minority sequence". The tree is rooted to the A/California/04/2009 virus (H1N1pdm09_Cali09; EPI_ISL_376192). Encircled tips denote the consensus majority sequence of the sample.

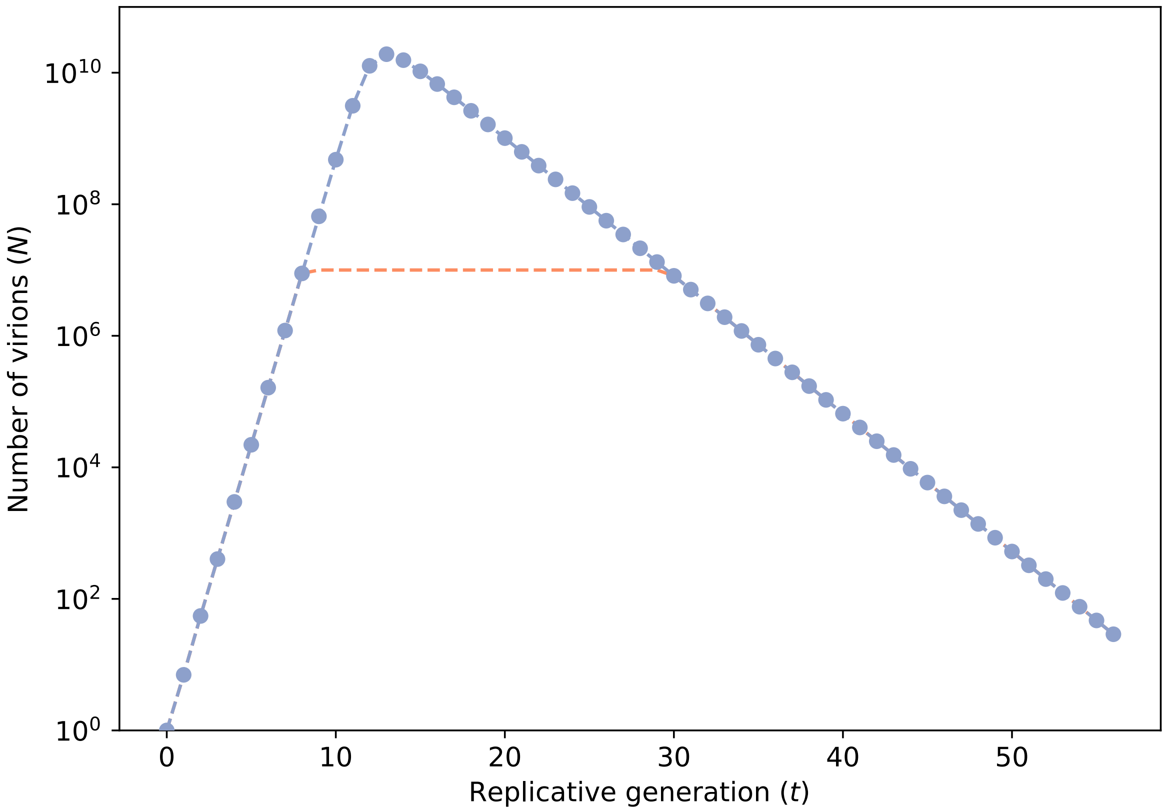


**Figure S11**: Number of virions ($N$) against replicative generation ($t$) based on a target cell-limited within-host model. Blue line with markers denotes the population size computed from the model. When $N > 10^7$ virions, we assumed that $N$ remained constant at $10^7$ (pink dashed line) to reduce computational costs of forward-time simulations.

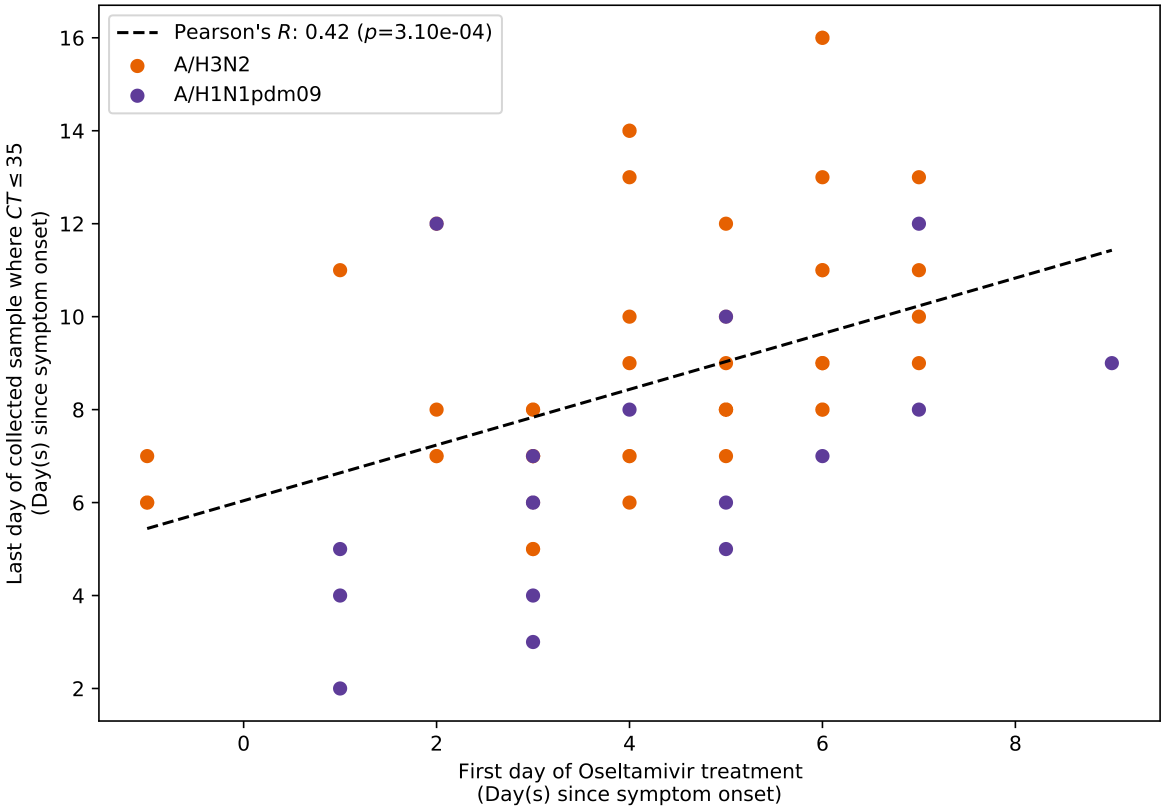


**Figure S12**: Pearson's correlation between the first day of Oseltamivir treatment administered to patients and the last day on which viral samples with cycle threshold (CT) values ≤ 35 were collected. Time is measured by number of days since symptom onset. Each point represents a patient included in this study who was treated with Oseltamivir (Table S4).

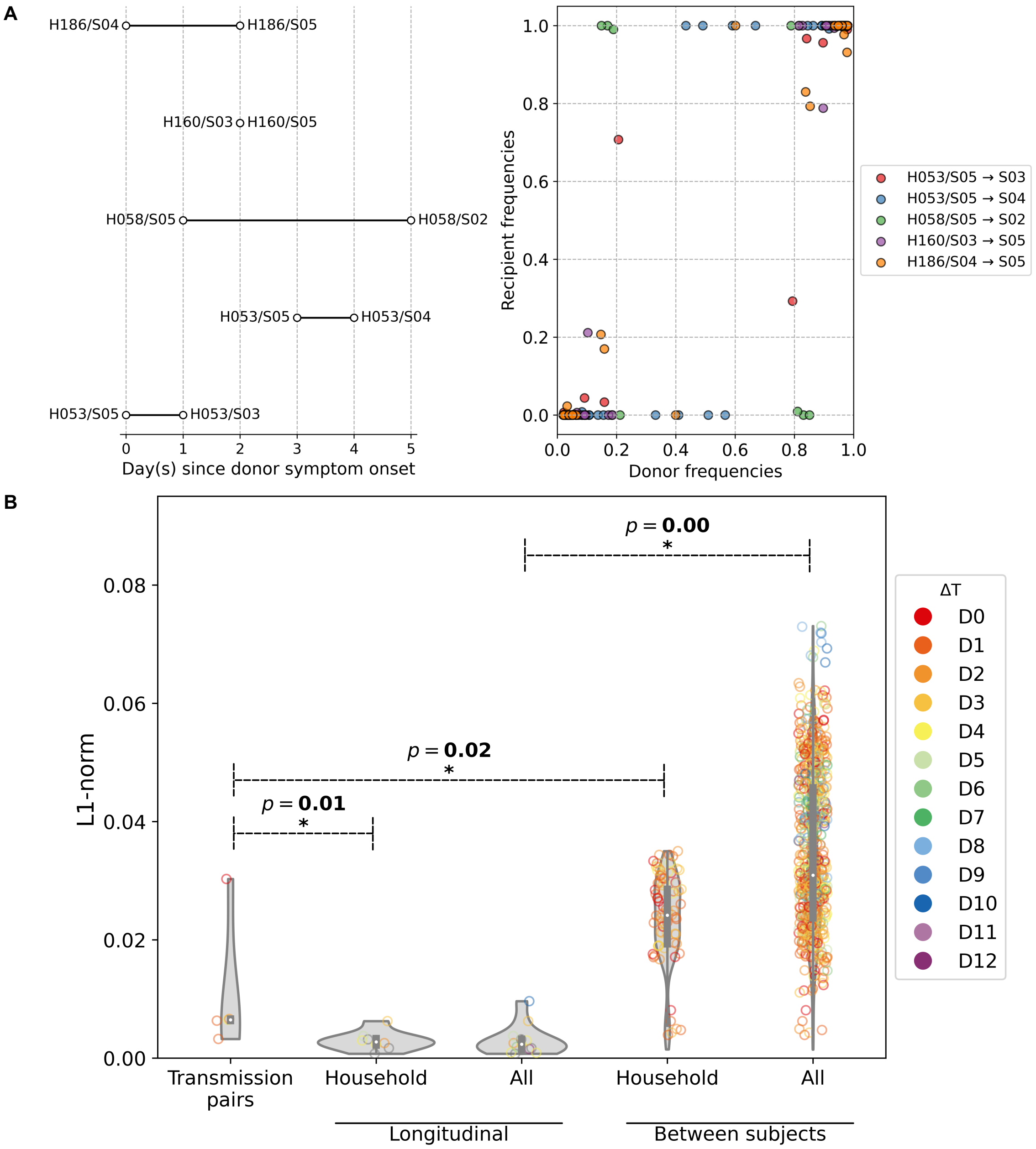


**Figure S13**: (**A**) Schematic of A/H1N1pdm09 virus household transmission pairs identified by epidemiological linkage and plotted based on timing of sample collection. (**B**) iSNV frequencies found in the donors and recipients of the five transmission pairs. (**C**) Violin plots of L1-norm pairwise genetic distance per site between different A/H1N1pdm09 virus sample pairs (each circle = 1 pair of virus samples). Transmission pairs are those represented in Figure 5A. Longitudinal pairs are made up of sample pairs collected from the same individual on the first and any other later timepoints on which the patient was sampled. These pairs are stratified by whether they were collected from households located in the same community (i.e. household) or combined with the rest of the analyzed A/H1N1pdm09 virus samples collected from hospitals (i.e. all). For the same aforementioned categories, we also plotted the distribution of L1-norm distances for pairs of viruses collected from different individuals. All circles are coloured by the difference in days between which the sample pairs were collected ($\Delta T$). All $p$-values reported are based on Mann-Whitney U tests which were used to determine if the L1-norm genetic distance distributions of the two categories marked by the ends of the horizontal line above are statistically distinct.

| Subtype | Gene | NS | S | Stop | NS/S |
|---|---|---|---|---|---|
| H3N2 | 1-PB2 | 4.78 | 5.59 | 0.04 | 0.85 |
| | 2-PB1 | 4.83 | 7.76 | 0.13 | 0.62 |
| | 3-PA | 4.92 | 9.77 | 0.19 | 0.50 |
| | 4-HA | 2.37 | 8.82 | 0.07 | 0.27 |
| | 5-NP | 6.90 | 4.12 | 0.06 | 1.68 |
| | 6-NA | 5.06 | 3.90 | 0.08 | 1.30 |
| | 7-M | 3.42 | 5.11 | 1.41 | 0.67 |
| | 8-NS | 5.75 | 10.04 | 1.10 | 0.57 |
| H1N1pdm09 | 1-PB2 | 4.98 | 3.99 | 0.12 | 1.25 |
| | 2-PB1 | 6.47 | 3.41 | 0.06 | 1.90 |
| | 3-PA | 5.26 | 4.13 | 0.15 | 1.27 |
| | 4-HA | 8.62 | 2.82 | 0.00 | 3.06 |
| | 5-NP | 4.92 | 4.07 | 0.11 | 1.21 |
| | 6-NA | 4.23 | 4.91 | 0.23 | 0.86 |
| | 7-M | 5.83 | 0.74 | 0.16 | 7.86 |
| | 8-NS | 6.53 | 3.61 | 0.00 | 1.81 |

**Table S1:** Mean number of nonsynonymous (NS), synonymous (S) and stop codon (Stop) variants per sample for each gene segment as well as the corresponding NS/S ratio.

| Subtype | Protein | Sample | Variant $\binom{i}{ii}$ | $LD$ | $LD'$ | $q_{10}$ | $q_{01}$ | $q_{11}$ |
|---|---|---|---|---|---|---|---|---|
| H1N1pdm09 | PB2 | 11-1015_D0 | *i.* S286G<br>*ii.* E343K | 0.04 | **1.00** | 0.00 | 0.00 | 0.04 |
| | HA | H160/S05_D7 | E261G<br>L455F | 0.04 | 0.30 | 0.34 | 0.08 | 0.15 |
| | NP | H053/S04_D8 | S314N<br>L358P | 0.08 | **1.00** | 0.00 | 0.02 | 0.09 |
| | NA | H160/S05_D6 | N386D<br>I389M | 0.10 | **1.00** | 0.00 | 0.00 | 0.12 |
| | M1 | H160/S05_D7 | L130P<br>G228D | 0.03 | **1.00** | 0.00 | 0.00 | 0.03 |
| | M2 | HCMC-1889_D0 | L46P<br>F48S | 0.16 | **1.00** | 0.00 | 0.04 | 0.21 |
| H3N2 | PB2 | 1219_D7 | I615V<br>S684F | 0.02 | **1.00** | 0.41 | 0.00 | 0.03 |
| | PB1 | 3611_D10 | K211R<br>E78K | 0.07 | **1.00** | 0.00 | 0.30 | 0.11 |
| | PA | 1224_D10 | E493G<br>F562S | 0.03 | **1.00** | 0.00 | 0.00 | 0.03 |
| | | 1691_D10 | K309R<br>R385I | 0.06 | **1.00** | 0.00 | 0.27 | 0.09 |
| | PA/PA-X | 3612_D7 | L175M<br>V201L | 0.07 | **1.00** | 0.00 | 0.27 | 0.11 |
| | NP | 1224_D10 | G384R<br>M426I | 0.05 | **1.00** | 0.34 | 0.00 | 0.08 |
| | | 1686_D8 | G384R<br>G102R | 0.08 | **1.00** | 0.00 | 0.25 | 0.13 |
| | | 1867_D7 | V197I<br>S353Y | 0.08 | **1.00** | 0.31 | 0.00 | 0.16 |
| | | 3615_D13 | G384R<br>A493T | 0.02 | **1.00** | 0.39 | 0.00 | 0.03 |
| | NS1 | 3615_D7 | R41G<br>E179G | 0.02 | **1.00** | 0.00 | 0.42 | 0.04 |
| | | 3615_D7 | H59L<br>E179G | 0.02 | **1.00** | 0.00 | 0.43 | 0.03 |
| | | 3611_D10 | V174A<br>T215S | 0.04 | **1.00** | 0.00 | 0.01 | 0.04 |
| | NEP | 3611_D10 | S22P<br>L63V | 0.04 | **1.00** | 0.00 | 0.01 | 0.04 |

**Table S2**: Potentially linked nonsynonymous variants in within-host A/H1N1pdm09 and A/H3N2 virus samples. Sample names are given in the format of "Patient ID_Days since symptom onset". Both linkage disequilibrium ($LD$) and the normalized $LD'$ measures are tabulated alongside the inferred maximum-likelihood haplotype frequencies ($q_{10}$ and $q_{01}$ are the haplotype frequencies with variant i or ii only while $q_{11}$ is the frequency of haplotypes encoding both variants).

**Table S3**: A/H3N2 segment-specific primers

| Gene | Amplicon | Sequence | Nucleotide coordinates Start | End |
|---|---|---|---|---|
| 1-PB2 | 1-F | AGCRAAAGCAGGTCAATTATATTCAG | -26 | 0 |
| | 1-R | GGTTCTGTCTAAGAATGTCCAC | 883 | 905 |
| | 2-F | ATTTCTCCCTTGATGGTTGCATAC | 592 | 616 |
| | 2-R | GACTCAGGACCGTTAATCTCC | 1611 | 1632 |
| | 3-F | GGTGTTTTCACAAGARGATTGC | 1206 | 1229 |
| | 3-R | AGTAGAAACAAGGTCGTTTTTAAAC | 2290 | 2315 |
| 2-PB1 | 1-F | AGCRAAAGCAGGCAAACCATTT | -23 | -1 |
| | 1-R | GCGATGCTCAGGATGTTTCTG | 999 | 1020 |
| | 2-F | ATGGTCACACAAAGAACAATAGG | 595 | 618 |
| | 2-R | CTGGTCCAAGGTCATTGTTTATC | 1602 | 1625 |
| | 3-F | AGCCCTGGRATGATGATGG | 1210 | 1229 |
| | 3-R | GACCAGTAGAAACAAGGCATT | 2301 | 2322 |
| 3-PA | 1-F | AGCRAAAGCAGGTACTGATTC | -23 | -2 |
| | 1-R | GTGACTTGGGTCTTCAATGCTC | 870 | 892 |
| | 2-F | GGGATTCCTTTCGTCAGTCC | 563 | 583 |
| | 2-R | CTCAAGGACACAGTATTTCTCC | 1611 | 1633 |
| | 3-F | GGCTCTTGGTGAAAACATGGC | 1104 | 1125 |
| | 3-R | GGATAACAAATAGTAGCACTGCC | 2154 | 2177 |
| 4-HA | 1-F | AGCRAAAGCAGGGGATAATTCTATTAA | -28 | -1 |
| | 1-R | CATTCCCTCCCAACCATTTTCTATG | 1062 | 1087 |
| | 2-F | CTCTATTGGGAGACCCTCAG | 254 | 274 |
| | 2-R | TCCTCAACATATTTCTCRAGGTCC | 1269 | 1293 |
| | 3-F | GACAATAGTAAAACCGGGAGAC | 750 | 772 |
| | 3-R | GACCAGTAGAAACAAGGGTG | 1717 | 1737 |
| 5-NP | 1-F | AGCRAAAGCAGGGTAGATAATCACTC | -44 | -18 |
| | 1-R | CTTCAAATGCAGCAGAATGGC | 998 | 1019 |
| | 2-F | ATGGATCCCAGAATGTGCTCTC | 475 | 497 |
| | 2-R | CAGTAGAAACAAGGGTATTTTTCC | 1499 | 1523 |
| | 3-F | TGAGGGAACTCGTCCTTTATG | 314 | 335 |
| | 3-R | TTATGGCCCAGTACCCGCTTC | 1145 | 1166 |
| 6-NA | 1-F | AGCRAAAGCAGGAGTAAAGATG | -19 | 3 |
| | 1-R | GTGTCTCCAACAAGTCCTG | 956 | 975 |
| | 2-F | GCATGGTCCAGCTCAAGTTG | 529 | 549 |
| | 2-R | GACCAGTAGAAACAAGGAGTT | 1431 | 1452 |
| | 3-F | GTGACAAGAGAACCTTATGTGTC | 346 | 369 |
| | 3-R | AGAAAATACCAGAATAACCGGACC | 1208 | 1232 |
| 7-M | 1-F | AGCRAAAGCAGGTAGATATTGAAAG | -24 | 1 |
| | 1-R | GACCAGTAGAAACAAGGTAG | 987 | 1007 |
| 8-NS | 1-F | AGCRAAAGCAGGGTGACAAAGAC | -25 | -2 |
| | 1-R | GACCAGTAGAAACAAGGGTG | 849 | 869 |

**Table S4**: Patients metadata (provided as an excel file).

**Table S5**: Acknowledgement table of reference sequences downloaded from GISAID.

| Isolate-ID | Isolate name | Country | Collection date | Originating Lab | Submitting Lab | Authors |
|---|---|---|---|---|---|---|
| EPI_ISL_103644 | A/BRISBANE/10/2007 | Australia | 2007-Feb-06 | Queensland Health Forensic and Scientific Services | WHO Collaborating Centre for Reference and Research on Influenza | Iannello,P; Komadina,N |
| EPI_ISL_376192 | A/California/04/2009 | United States | 2009-Jan-01 | | Import from public-domain | Dpcc,C.D. |
